# NEIL2 regulates RNA polymerase II dynamics to suppress transcription–replication conflicts in pancreatic cancer

**DOI:** 10.64898/2026.10.19.760262

**Authors:** Fan Meng, Anthony Fernandez, Changfei Luan, Marc A. Attiyeh, Qianhua Feng, Alexander Lee, Haihua Feng, Tiane Li, Tingting Zhou, Yun R Li, Russell Rockne, Terence Williams, David Dawson, Binghui Shen, Jen Jen Yeh, Yilun Liu, Maria Ninova, Mustafa Raoof

## Abstract

Transcription–replication conflicts (TRCs) arise from spatiotemporal interference between replication and transcription machinery and contribute to genome instability in cancer cells. In pancreatic ductal adenocarcinoma (PDAC) cells, oncogenic KRAS drives TRC formation, and base excision repair (BER) contributes to TRC resolution as an adaptive response. Here, we show that TRCs are enriched in pancreatic tumor sections from patients and are associated with malignant progression. We identify NEIL2, a bifunctional DNA glycosylase in the BER pathway, as a critical regulator of TRCs in PDAC cells. Mechanistically, NEIL2 loss causes aberrant RNA polymerase II (RNAPII) occupancy at transcription start sites and downstream of transcription end sites. Dysregulated RNAPII dynamics promote R-loop-dependent and -independent TRCs contributing to replication stress, DNA damage, and impaired cancer cell proliferation. Notably, the DNA-binding motif of NEIL2 necessary for canonical BER function is dispensable for TRC regulation. Together, these findings reveal an unrecognized role for NEIL2 in regulating RNAPII dynamics and mitigating TRCs in pancreatic cancer cells.

## INTRODUCTION

Transcription-replication conflicts(TRCs) arise from collisions between the transcription and replication machinery and represent a major endogenous source of DNA damage (*1, 2*). Head-on TRCs preferentially activate the Ataxia Telangiectasia and Rad3-related (ATR) pathway, whereas co-directional conflicts trigger the Ataxia-Telangiectasia Mutated (ATM) pathway (*3*). Unresolved TRCs induce severe genomic instability, including double-strand breaks (DSBs) (*4–6*). Our previous study demonstrated that base excision repair (BER), which protects against oxidative DNA damage, also plays a critical role in resolving oncogenic KRAS-driven TRCs. Consequently, inhibition of BER at abasic(AP) site processing step increases TRC and promotes the accumulation of TRC-related DNA damage (*7*). However, the specific BER components involved in TRC regulation and their underlying mechanism remained unclear.

Pancreatic ductal adenocarcinoma (PDAC) is the third leading cause of cancer-related deaths in the US and the sixth leading cause worldwide. Systemic therapy is the only option in 75% of patients who are present with metastatic disease, with a 5-year survival rate of 13% (*8*). Approximately 90% of PDACs harbor oncogenic KRAS mutations and mutant-KRAS was thought to be undruggable until recently (*9*). Although clinical benefit of novel mutant-allele-specific and pan-KRAS inhibitors (*10–13*) has been demonstrated (*13*), therapeutic resistance through RAS-bypass is inevitable (*9, 14–16*). Thus, alternative therapeutic strategies are urgently needed. Exploiting cancer-associated genomic instability has emerged as a promising therapeutic strategy in PDAC(*17*). Consistent with this concept, we previously demonstrated that exacerbating oncogene-driven TRCs provides therapeutic benefit(*18*). We therefore hypothesized that that identify specific BER factor involved in TRC regulation would reveal selective therapeutic vulnerabilities in PDAC.

Using siRNA screens on human pancreatic cancer cells, we identified NEIL2 (Nei Like DNA Glycosylase 2), a bifunctional DNA glycosylase in the Fpg/Nei family, as a cancer-selective BER component that reduces TRCs (*7, 19*). Unlike other glycosylases, NEIL2 functions are non-redundant. It preferentially targets lesions in actively transcribed genes through transcription-coupled BER (TC-BER) and functionally associates with RNA polymerase II (RNAPII) via recruitment of TC-BER factors, such as excision repair cross-complementation group 6 (ERCC6) (*20–22*). Importantly, loss of NEIL2 in normal cells has non-lethal effects. For instance, NEIL2-deficient mice, despite accumulating oxidative DNA lesions, do not exhibit overt developmental or physiological abnormalities(*23*).In contrast, deletion of essential downstream BER factors, including Apurinic/apyrimidinic endonuclease 1(APE1), DNA polymerase β (Pol β), and DNA ligase IIIα (LIG3), leads to embryonic lethality(*23*). Together with our siRNA screening results, these observations suggest that NEIL2 loss is well tolerated in normal tissue, whereas NEIL2 is essential for TRC resolution in PDAC cells.

Here, we establish the clinical relevance of TRCs in PDAC. Proximity-ligation assay (PLA)-based detection in patient samples, showed that TRCs are enriched in tumor cells compared to stroma and adjacent normal ductal epithelium, and positively correlate with tumor size and grade. We further validate NEIL2 as a critical regulator of TRCs using CRISPR-mediated gene knockout and show that this is independent of its DNA binding domain. Loss of NEIL2 results in aberrant RNAPII occupancy at transcription start (TSS) and end sites (TES) promoting TRC formation and DNA damage and impairs PDAC cell viability and proliferation in 2D and 3D *in-vitro* models, without affecting normal cells. Overall, we define a non-canonical and a critical adaptive role of NEIL2 in mitigating TRCs in PDAC.

## RESULTS

### Human PDAC shows enrichment of TRCs

We optimized the PLA method to determine the extent of TRC accumulation in formalin-fixed paraffin embedded human cancer tissue. We performed RNAPII-PCNA PLA in primary PDAC tumor and paired normal tissue derived from 22 treatment-naive patients undergoing curative-intent pancreatectomy.

CK19 (green) was used as an epithelial tumor marker on both tumor sections and matched normal tissues to distinguish between cancer and non-cancer-cell components (*24, 25*). CK19-positive (CK19(+)) cancer cells exhibited two-fold increase in TRC-PLA foci compared with CK19-negative (CK19(-)) components of the tumor microenvironment, (*P* < 0.0005). In contrast, TRC-PLA signals were largely undetectable in normal tissues, including CK19(+) ductal epithelial cells (Fig. 1A and Fig. S1A).

**Figure 1:**
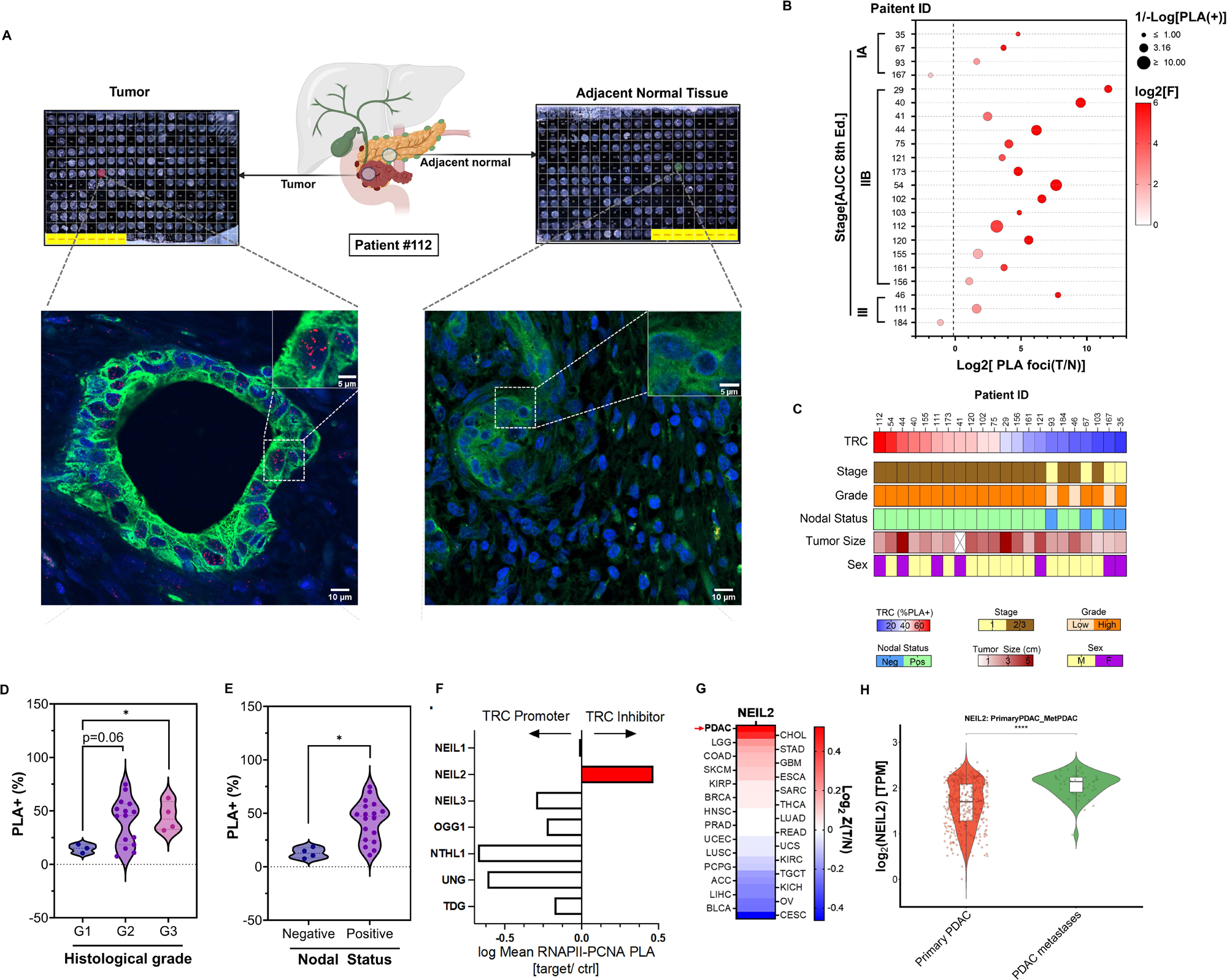
Human PDAC shows enrichment of TRCs. **(A)** Representative RNAPII–PCNA proximity ligation assay (PLA) images of pancreatic ductal adenocarcinoma (PDAC) tumor tissues and matched adjacent normal pancreatic tissues from patient # 112. Low-magnification tissue overview and high-magnification images are shown. Red staining indicates RNAPII–PCNA PLA foci, green staining indicates CK19, and blue staining shows nuclei counterstained with DAPI. **(B)** Quantification of RNAPII–PCNA PLA signals in PDAC specimens from individual patients. The x-axis shows the log₂-transformed enrichment of PLA signals in tumor tissues relative to matched adjacent normal tissues [log₂(T/N)]. Bubble size represents the inverse log-transformed frequency of PLA-positive cells [1/−log(PLA⁺%)]. Bubble color indicates the statistical significance of PLA enrichment, represented as the log₂-transformed *F* statistic derived from the comparison between tumor and matched adjacent tissues. Each bubble represents an individual patient sample. **(C)** Clinical annotation heatmap summarizing clinicopathological characteristics of the PDAC patient cohort. Patient sex is indicated by color (yellow, male; purple, female). Histological grade is represented by a color gradient from light to dark corresponding to Grade 1–3 tumors. Nodal -status is represented by a color(blue, negative; purple, positive). Tumor stage is represented by a color gradient from light to dark corresponding to Stage I–III disease. Additional annotations include tumor size, percentage of PLA(+) cells as determined by RNAPII–PCNA PLA analysis. (**D**) Comparison of TRC burden between nodal status-negative and nodal status-positive tumors **(E)** The relative frequency of tumors is plotted according to the percentage of PLA-positive tumor cells using 10-percentage-point bins, with values shown at each bin center. The cohort median of 25% PLA-positive cells (dotted line) was used as the cutoff to stratify patients into PLA-low (blue) and PLA-high (green) groups for survival analysis. PLA-positive cells were defined as tumor cells containing >3 PLA foci per nucleus. (**F**)RNAPII–PCNA PLA-based screening of BER glycosylases using a DDR-focused siRNA library. Candidate genes were ranked according to their effects on TRC levels following siRNA-mediated depletion. **(G)** Pan-cancer analysis of NEIL2 expression across TCGA solid tumor cohorts. Heatmap shows differential NEIL2 expression between primary tumors from TCGA and corresponding normal tissues from GTEx. Color intensity represents relative expression levels, with warmer colors indicating higher expression and cooler colors indicating lower expression relative to normal tissues. **(H)** NEIL2 expression in primary and metastatic PDAC from UNC cohort. Differences were assessed using the Wilcoxon rank-sum test. \*\*\**P* < 0.0001.

We further correlated TRC burden with histologic grade, lymph node status and tumor size (Fig. 1C). TRC-PLA foci in cancer cells positively correlated with tumor size (Fig. S1B). In addition, the mean proportion of PLA(+) cells (defined as 3+ foci) was approximately three-fold higher in CK19(+) malignant cells from grade 2 (*P* = 0.06) and grade 3 (*P* = 0.02) tumors compared to grade 1 tumors (Fig. 1D). Furthermore, TRC burden was significantly associated with lymph node involvement, with node-positive (N1/N2) tumors exhibiting a higher proportion of PLA-positive cells than node-negative (N0) tumors (Fig. 1E).

To evaluate whether TRC levels correlate with disease prognosis, we first examined the frequency distribution of PLA(+) tumor-cell percentages across the patient cohort. This analysis demonstrated a bimodal distribution. The bottom tercile was categorized as PLA-low and middle and top tercile was categorized as PLA-high (Fig. S1C left panel). Kaplan–Meier survival analysis demonstrated that the PLA-low group had a more than two-fold higher estimated 5-year disease-free survival (DFS) probability than the PLA-high group (62.5% versus 25.4%), although the difference in DFS did not reach statistical significance (log-rank P = 0.08) (Fig. S1C right panel). Collectively, these data demonstrate that TRCs are enriched in human PDAC cells, correlate with adverse pathological features, and may be associated with poor prognosis.

We have previously identified BER pathway as a key regulator of TRCs in PDAC(*7*). Comparing the effects of BER glycosylases on TRCs in the Panc1 cell line by siRNA knockdown, NEIL2 was the only BER glycosylase contributing to TRC mitigation (Fig. 1F). Interestingly, comparison of NEIL2 expression between TCGA tumor datasets and GTEx normal tissue datasets via GEPIA2 (*26*) revealed that the relative NEIL2 tumor-to-normal (T/N) expression ratio was highest in PDAC among 28 solid tumor types (Fig. 1G). Meanwhile, analysis of single cell transcriptomic profile in normal pancreas tissue using the Human Protein Atlas database (*27*) showed extremely low NEIL2 expression across all normal cell types (Fig S1D), suggesting the NEIL2 upregulation is a cancer-selective dependency in PDAC. To further evaluate the clinical relevance of NEIL2 in PDAC, we interrogated the UNC database. Bulk RNA-sequencing analysis of primary and metastatic PDAC samples showed significantly higher NEIL2 expression in metastatic lesions than in primary tumors (*P* < 0.0001; Fig. 1H), suggesting a strong association between NEIL2 expression and metastatic phenotype regardless of transcriptomic sub-type (basal-like versus classical Fig. S1E). Taken together, these findings suggest a model in which NEIL2 expression represents a cancer-associated adaptive response to TRC accumulation in PDAC and is associated with aggressive disease phenotypes.

### NEIL2 deprivation leads to TRC accumulation and DNA damage in PDAC cells

To functionally validate NEIL2 as a regulator of TRCs in PDAC, we generated NEIL2-knockout PANC-1 cells(N2KO) using CRISPR–Cas9. N2KO cells exhibited significantly increased TRC levels compared with Control cells expressing a scrambled sgRNA (Fig. 2A). To determine the consequences of NEIL2 loss for DNA replication dynamics, we performed DNA fiber assays and observed increased replication fork stalling and reduced fork progression rates in N2KO cells relative to controls (Fig. 2B). Consistent with elevated TRCs and replication stress, N2KO cells exhibited activation of the ATR–Chk1 signaling axis and increased nuclear γH2AX levels, indicating enhanced DNA damage (Fig. 2C). Using a cell-cycle synchronization and release strategy, we further found that replication stress induced by NEIL2 deficiency impaired S-phase progression (Fig. 2D). Notably, immunofluorescence staining revealed a pan-nuclear γH2AX pattern in N2KO cells (Fig. 2E), consistent with widespread replication stress. Collectively, these findings demonstrate that loss of NEIL2 leads to TRC accumulation and widespread replication stress, thereby impairing DNA replication and activating the DNA damage response in PDAC cells.

**Figure 2:**
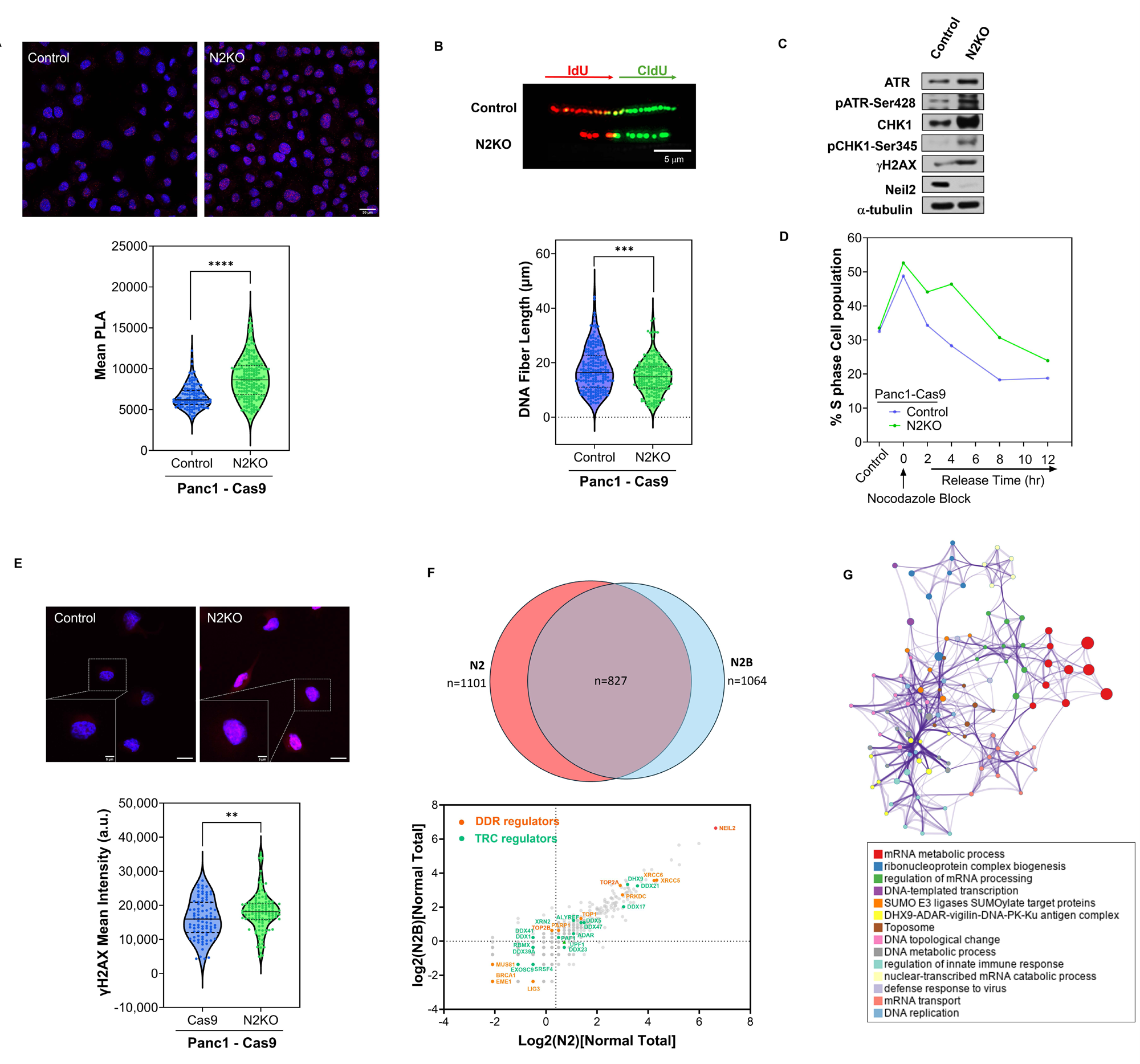
NEIL2 deprivation leads to TRC accumulation and DNA damage in PDAC cells. **(A)** Representative RNAPII–PCNA proximity ligation assay (PLA) images in Panc1-Cas9 control(Control) and NEIL2-knockout (N2KO) cells. Quantification of PLA signal intensity demonstrates increased TRC accumulation following NEIL2 depletion. Each dot represents an individual nucleus. Scale bar, 30 μm. **(B)** DNA fiber assay assessing replication fork progression in Control and N2KO cells. Actively replicating DNA was sequentially labeled with ldU (red) followed by CldU (green) to monitor replication fork dynamics. Representative DNA fiber images are shown above. Quantification of total replication track length revealed reduced fork progression in NEIL2-deficient cells, indicating increased replication stress and impaired DNA replication. **(C**) Western blot of replication stress and DNA damage signaling pathways in Control and N2KO cells. Protein levels of ATR, phospho-ATR (Ser428), CHK1, phospho-CHK1 (Ser345), γH2AX, and NEIL2 are shown. α-tubulin serves as loading control. **(D)** Representative cell-cycle progression analysis following nocodazole synchronization and release in Control and N2KO cells. Cells were synchronized by nocodazole treatment and released into fresh medium for the indicated time intervals. The percentage of S-phase cells was quantified at each time point by flow cytometry. **(E)** Representative immunofluorescence images of γH2AX in Control and N2KO cells. γH2AX foci are shown in red, and nuclei are counterstained with DAPI (blue). Quantification of nuclear γH2AX signal intensity is presented as violin plots, with each data point representing an individual nucleus. Scale bar, 15 μm **(F)** Identification of NEIL2-associated TRC regulatory networks. Venn diagram showing overlap between proteins identified in NEIL2 immunoprecipitates from N2 and N2B. Scatter plot displays normalized total peptide count of NEIL2-associated proteins, highlighting known DNA damage response (DDR) and transcription–replication conflict (TRC) regulators. **(G)** Functional enrichment analysis of the NEIL2 interactome. Network visualization of enriched biological functions identified among NEIL2-associated proteins involved in TRC regulation. Node size corresponds to the number of proteins associated with each term, and node color indicates functional cluster membership. Edges connect terms with similarity scores >0.3, with edge thickness proportional to similarity. The network was generated in Cytoscape using a force-directed layout with edge bundling. One representative term from each cluster is labeled.

Next, we investigated the mechanism by which NEIL2 regulates TRCs. We performed co-immunoprecipitation coupled with mass spectrometry (Co-IP/MS) using Flag-tagged NEIL2 transiently expressed in Panc-1 cells. We analyzed two independently prepared lysates in parallel. To improve recovery of chromatin-associated protein complexes, one lysate was additionally treated with benzonase – a genetically engineered endonuclease - which we designated as the N2B group. Despite the different nuclease conditions, the two datasets showed substantial overlap, with 827 proteins reproducibly detected in both preparations, representing 75.1% and 77.7% of the candidate proteins identified in N2 and N2B, respectively. These results indicate that the NEIL2-associated proteome was highly reproducible across the two experimental conditions.

As expected, the NEIL2 interactome includes several well-known DNA damage response (DDR) factors, including XRCC5/XRCC6 (Ku80/Ku70) and PRKDC (DNA-PKcs, key mediators of double-strand break repair (*28*) (Figure 2F, lower panel). We also identified TOP1, TOP2A, and TOP2B (*29–31*), which alleviate transcription-associated torsional stress, as well as PARP1 and LIG3 (*21, 32*), that are established downstream components of the BER pathway (Fig. 2F, lower panel). Furthermore, NEIL2 interactome includes multiple RNA helicases implicated in transcription and R-loop metabolism, including DDX5 (*33*), DDX21 (*34*), DDX17 (*35*), and DHX9 (*36*). Altogether, 23 previously reported TRC regulators were reproducibly detected in both IP groups (Fig. S2A). Metascape analysis of these factors revealed a functional network connecting DDR, TRC regulation, and transcription-associated processes, mRNA metabolism, mRNA processing, and RNA processing, as well as DDR-related pathways such as DNA topology regulation, DNA metabolic processes, and the DNA-PK–Ku complex (Fig. 2G). Indeed, several factors in this network, including TOP1 (*37*), TOP2A (*38*), XRCC5 (*39*), and PARP1 (*40*) link the TRC-regulatory network with the DNA topology and repair factors associated with NEIL2, suggesting a functional connection between NEIL2-mediated DNA repair and TRC regulation. Importantly, similar functional themes were independently identified through unbiased GO analysis of high-confidence NEIL2 interactors, which also revealed enrichment of RNA- and transcription-associated processes, as well as DNA replication and double-strand break repair pathways (Fig. S2B).

Collectively, these findings suggest that NEIL2 beyond its canonical BER functions, participates broadly in TRC regulatory networks through the coordination of transcription-associated DNA repair, resolution of topological stress, R-loop metabolism, and processing of TRC-induced DNA lesions. Interestingly, comparison of the NEIL2 interactomes from Panc1 and normal pancreas cell line, HPNE, revealed only 70 shared interactors. Most established TRC regulators were selectively enriched in Panc1 cells, whereas only UPF1 (*41*) and DDX21 were common to both datasets, and SRSF2 (*42*) and MCM5 (*43*) were uniquely identified in HPNE cells (Fig. S2C). These findings suggest potential differences in the NEIL2-associated TRC regulatory networks between PDAC and nonmalignant pancreatic cells.

### NEIL2 resolves TRCs via TC-BER pathway but independent of DNA binding

Given the role of NEIL2 in TC-BER and TRCs (*7, 21*), we investigated whether NEIL2 resolves TRCs specifically through the TC-BER pathway. Because the BER pathway is a primary defense against oxidative DNA damage(*44–46*), we asked whether elevated TRCs following loss of NEIL2 are due to impaired resolution of oxidative stress. To address this, we performed flow cytometric analysis of ethanol-fixed Panc1-Cas9 Control and N2KO cells co-stained with DAPI and 8-oxo-dG - a marker of oxidative DNA lesions(*20*). In both cell lines, oxidative lesions progressively accumulated as cells advanced through the cell cycle. However, N2KO cells exhibited significantly higher levels of 8-oxo-dG in G1, S and G2 phases compared with controls (Fig. 3A and S3A), indicating enhanced accumulation of oxidative stress–induced DNA damage. To determine whether the oxidative DNA damage phenotype associated with NEIL2 loss is reflected in patient tumors, we analyzed whole-genome mutation and matched expression data from 254 pancreatic cancers in the ICGC cohort. NEIL2 expression inversely correlated with the fraction of C>A transversions, a mutational signature of unrepaired 8-oxoguanine (Spearman r = −0.18, *P* = 0.004). Consistently, NEIL2-low tumors exhibited a higher C>A mutation fraction than NEIL2-high tumors (two-sided Mann–Whitney *P* = 0.036; Fig. S3B), supporting increased 8-oxo-dG accumulation following loss of NEIL2 in vitro (Fig. S3A). To determine whether this oxidative stress contributes to TRCs and replication stress, we treated both Panc1-Cas9 Control and N2KO cells with the antioxidant N-acetylcysteine amide (NACA). Antioxidant treatment markedly reduced TRC levels in both cell lines, restoring them to near-basal levels comparable to untreated Cas9 controls (Fig. 3B). In parallel, NACA rescued replication fork progression as assessed by DNA fiber analysis (Fig. S3C). These results demonstrate that oxidative stress is a major contributor to TRC accumulation and replication stress in NEIL2-deficient cells.

**Figure 3:**
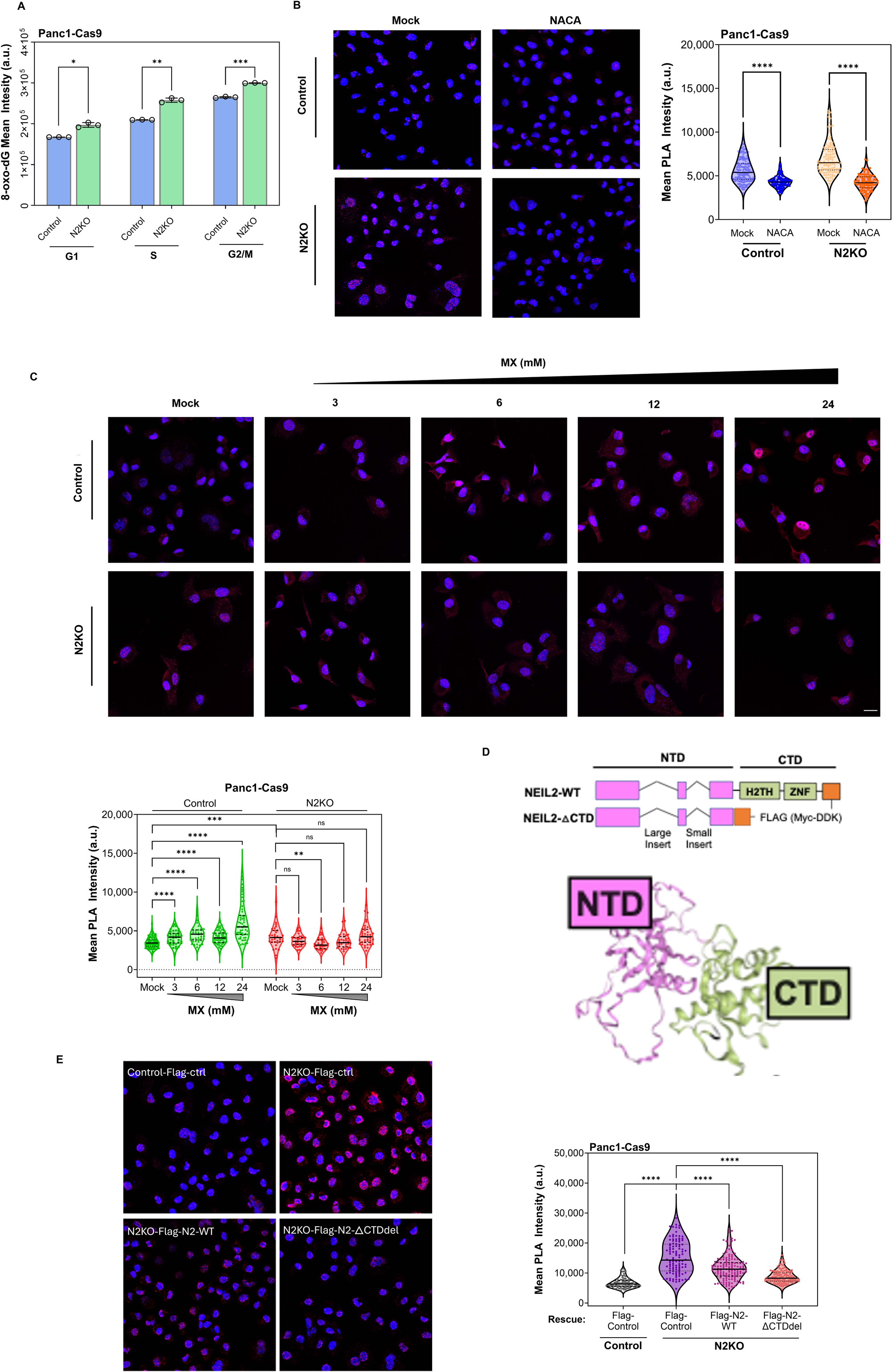
NEIL2 resolves TRCs via TC-BER pathway but independent of DNA binding. **(A)** Flow cytometric quantification of 8-oxo-dG levels during different cell-cycle phases in Panc1-Cas9 control(Control) and Panc-1 N2KO(N2KO) cells. Each dot represents the mean 8-oxo-dG fluorescence intensity from one independent biological replicate (n = 3). **(B)** RNAPII–PCNA PLA analysis following treatment with 2 mM N-acetylcysteine amide (NACA) for 2 hours. Representative images show PLA foci (red) and DAPI-stained nuclei (blue). Scale bar, 30 μm. **(C)** RNAPII–PCNA PLA analysis in Control and N2KO cells treated with increasing concentrations of methoxyamine (MX) for 2 hours. Representative images show PLA foci (red) and DAPI-stained nuclei (blue). Scale bar, 30 μm. **(D)** Schematic representation of wild-type (WT) and C-terminal deletion (ΔCTD) NEIL2 constructs used for functional rescue experiments. **(E)** RNAPII–PCNA proximity ligation assay (PLA) functional rescue experiments of Control and N2KO cells transfected with the indicated constructs are shown. PLA signals are displayed in red and nuclei are counterstained with DAPI (blue). Scale bar, 30 μm.

To test if BER pathway contributes to NEIL2-TRC regulation, we treated Panc1-Cas9 Control and N2KO cells with increasing concentrations (3–24 mM) of the BER pathway inhibitor methoxyamine (MX)(*47*). MX reacts with abasic sites resulting from BER glycosylase active form a bulky lesion that interferes with the critical BER function of Apurinic/apyrimidinic endonuclease 1 (APE1). In Control cells, MX induced TRCs in a clear dose-dependent manner, even at sublethal concentrations (3–6 mM). N2KO cells, on other hand, were largely insensitive to MX treatment, even at high concentrations (12–24 mM), showing no additive increase in TRCs (Fig. 3C). These findings suggest that NEIL2 deficiency nearly saturates BER inhibition–induced TRC formation, supporting an essential role of the NEIL2-BER axis in TRCs regulation.

Given the role of NEIL2-BER axis in TRC regulation, we next sought to assess whether the glycosylase activity of NEIL2 is required for interactions with TRC and R-loop regulators. We constructed full-length NEIL2 (NEIL2-WT) and a C-terminally truncated mutant (NEIL2-CTDΔ) (Fig.3D), which lacks DNA-binding capacity and attenuates the catalytic activity of NEIL2 but retains interaction with TC-BER components(*48*) and the RNAPII complex (Fig S3E). Transient expression of either WT or CTDΔ construct partially rescued the elevated TRCs observed in NEIL2-knockout cells (Fig. 3E). Collectively, these findings suggest that NEIL2-mediated regulation of TRCs is independent of its DNA-binding, pointing to additional mechanistic roles of NEIL2 in maintaining transcription–replication homeostasis.

### RNAPII dysregulation at TSS and TES drives TRCs in NEIL2-knockdown cells

Endogenous oxidative stress in N2KO cells promotes replication fork stalling and TRCs. We next asked whether replication stress itself is the direct driver of TRC formation in the absence of NEIL2. To test this, we applied a short pulse (2 hrs) of the replication stress–inducing agent aphidicolin (APH) to transiently arrest global replication fork progression(*49*) in both Panc1-Cas9 Control and N2KO cells (Fig. 4A). Replication arrest was confirmed in both cell lines (Fig. S3C). In parallel, we monitored RNAPII dynamics, TRCs and DNA damage responses over time after APH was removed (Fig. 4B and Fig. S4A). In Control cells, APH treatment rapidly increased TRCs and γH2AX (Fig. 4A,B), consistent with transcription machinery encountering stalled replication forks(*50*). Following APH removal, both of TRC and γH2AX signals declined in control cells, consistent with recovery from replication stress (Fig. S4A). In contrast, the initial APH pulse did not further increase TRC or γH2AX signals in N2KO cells(Fig. 4A,B). During recovery, TRC signals declined in N2KO cells while γH2AX remained elevated (Fig. S4A). Notably, at 24 hours after APH removal, TRCs had returned to approximately baseline levels, but γH2AX remained substantially higher in N2KO cells than in control cells. Overall, these results suggest that replication fork stalling alone does not drive the elevated TRC formation in N2KO cell, although global replication stress can exacerbate genotoxic stress.

**Figure 4:**
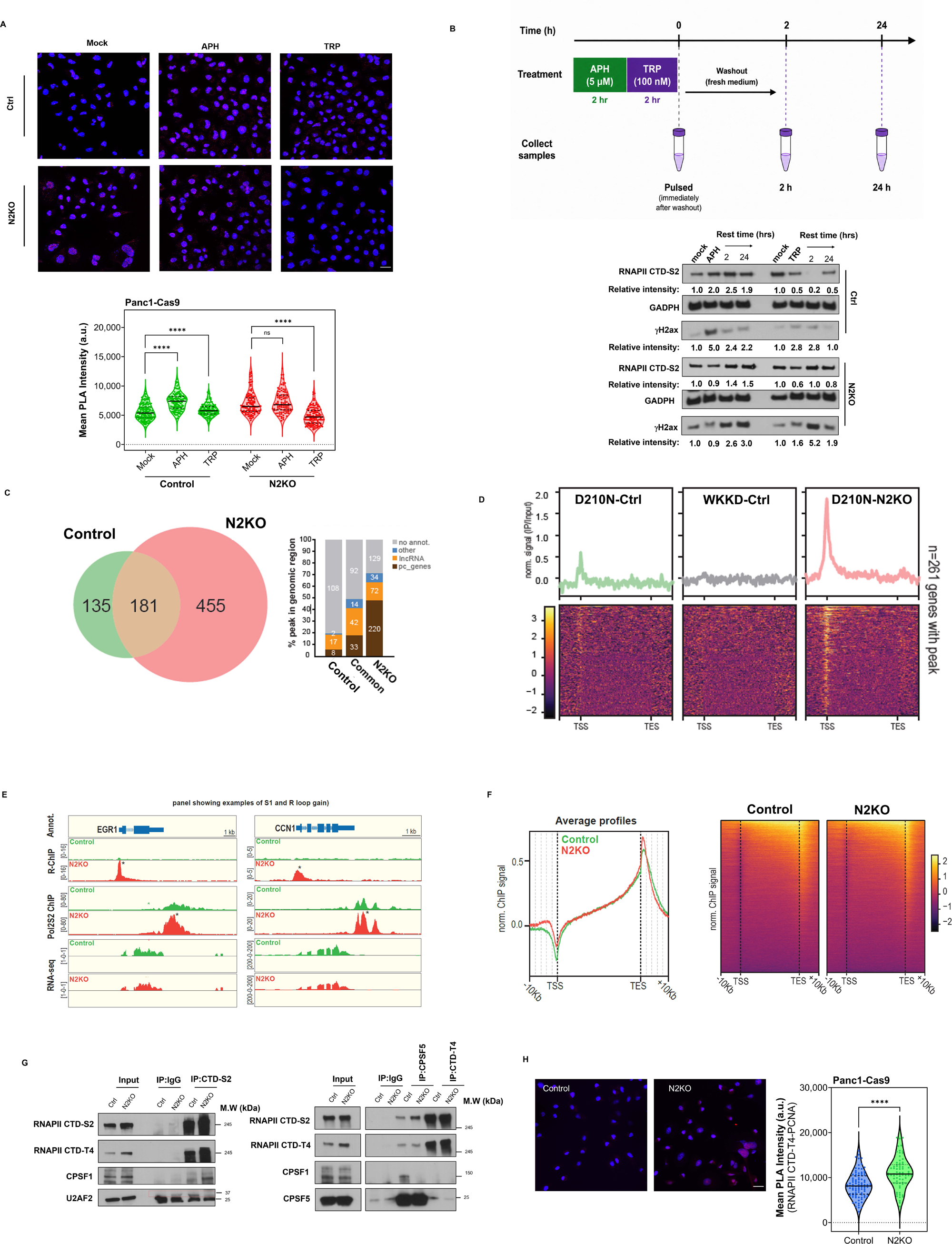
RNAPII dysregulation at TSS and TES drives TRCs in NEIL2-knockdown cells. **(A)** Proximity ligation assay (PLA) analysis following treatment with 5mM aphidicolin (APH) and 100 nM triptolide (TRP) for 30 mins. Representative images and quantification demonstrate that replication stress and transcription-initiate inhibition differentially affect TRC accumulation in Control and N2KO cells. Scale bar, 30 μm. **(B)** Schemic workflow of pulse-and-recovery experiment with APH and TRP treatment respectively. Immunoblot analysis of RNA polymerase II CTD Ser2 phosphorylation (RNAPII CTD-S2) and γH2AX following transient aphidicolin (APH) or triptolide (TRP) treatment and recovery as indicated time. Relative protein intensities are indicated below each blot and were quantified by densitometric analysis, normalized to GAPDH, and expressed relative to mock-treated samples (set to 1). **(C)** Venn diagram illustrating the overlap between genes with increased R-loop accumulation and altered transcriptional activity in N2KO cells. Stacked bar plots show the genomic distribution of differential R-loop peaks that are unique to Control cells, unique to N2KO cells, or shared between both conditions. Genomic annotations were assigned based on peak localization relative to gene features, revealing preferential accumulation of N2KO-associated R-loops at transcriptionally active genomic regions. **(D)** Metagene profiles and heatmaps of R-ChIP-seq signals in Panc1 cells expressing catalytically inactive RNase H1 (D210N), RNase H1-WKKD negative control, or D210N in N2KO cells. Genome-wide analysis of 261 R-loop-positive genes demonstrated elevated R-loop accumulation upon NEIL2 depletion, particularly at promoter-proximal regions and transcription start sites (TSSs). Metagene plots represent average normalized R-ChIP signal distribution, and heatmaps show normalized signal intensity across individual genes ranked by R-loop enrichment. **(E)** Genome browser tracks of representative loci (EGR1 and CCN1) showing R-loop accumulation and elongating RNA polymerase II occupancy. **(F)** Aggregate profiles and heatmaps of elongating RNA polymerase II (POLR2S2) ChIP-seq signals across transcription units in Control and N2KO cells. **(G)** Co-immunoprecipitation analysis of elongating RNA polymerase II (RNAPII CTD-S2) and termination-associated RNA polymerase II (RNAPII CTD-T4) complexes with key 3′-end RNA processing factors, including CPSF1, CPSF5, and U2AF2, in Control and N2KO cells. **(H)** RNAPII CTD-T4–PCNA PLA analysis in Control and N2KO cells. Representative images show PLA signals (red) and DAPI-stained nuclei (blue) and quantification of PLA signal intensity. Scale bar, 30 μm.

To test the transcription dependence of NEIL2-associated TRCs, we treated cells with a 2-hour pulse of triptolide (TRP). TRP inhibits transcription initiation by targeting the XPB subunit of TFIIH, thereby blocking promoter opening and promoting turnover of promoter-associated RNAPII without affecting DNA replication (Fig. S3C) or elongating RNA polymerases(*51, 52*). TRCs in N2KO were rapidly reduced by a short 2-hour pulse of TRP (Fig. 4A). In contrast, Control cells exhibited mildly increase in TRCs with TRP pulse (Fig. 4A). The differential response to TRP suggests that TRCs in N2KO cells preferentially involve RNAPII complexes at initiation or transcriptionally engaged RNAPII complexes sensitive to TRP. To explore the basis of this divergent TRC response, we next examined the RNAPII dynamics under TRP pulse and washout. Although the initial TRP pulse caused comparable reduction in the elongating RNAPII(CTD-S2) signal in both control and N2KO, their response diverged during washout (Fig. 4B). In control cell, RNAPII CTD-S2 was depleted 2 hours after TRP removal, potentially reflecting clearness of RNAPII complex following transcription completion (Fig. 4B). In N2KO cell, the RNAPII CTD-S2 returned to baseline levels within 2 h after TRP washout, suggesting rapid recovery of stalled RNAPII or impaired RNAPII clearance in N2KO cells. Recovery of CTD-S2 coincided with elevated γH2AX levels without further TRC formation (Fig. 4B and S4B), suggesting that prolonged RNAPII retention on chromatin in N2KO cells during TRP washout leads to TRC-independent DNA damage. Together, these findings suggest that NEIL2 contributes to TRC homeostasis by regulating RNAPII occupancy and dynamics during transcription.

To investigate whether NEIL2 contribute to TRC resolution beyond its canonical function in BER, we focused on NEIL2-CTDΔ interacting proteome as it rescued the N2KO TRC phenotype partially. Co-IP mass spectrometry results showed that NEIL2-CTDΔ retained 116 out of 182 proteins enriched with NEIL2-CTDΔ were also detected with NEIL2-WT (Fig S3D). The common hits from these two constructs were associated with RNA processing factors and R-loop regulators.

Since head-on TRCs generate R-loops (*3*), we assessed global R-loop abundance by dot blotting genomic DNA from Panc1-Cas9 Control and N2KO cells using the S9.6 antibody. N2KO consistently resulted in elevated R-loop levels compared with Control cells (Fig. S4C), indicating that NEIL2 suppresses endogenous R-loop accumulation. We also performed genome wide mapping of R-loops using R-ChIP sequencing. In this assay, catalytically inactive RNase H1 (D210N) was used to capture R-loops, while the catalytic and nucleic acid–binding–deficient RNase H1 mutant (WKKD) served as a negative control (*53*). Consistent with dot-blot results, control cells display relatively small number of R-loop peaks, most of which fell outside annotated protein-coding and lncRNA genes, the major classes of RNA polymerase II-transcribed loci. In contrast, N2KO cells showed elevated R-loops with 455 unique R-loop peaks (Fig. 4C), the majority of which mapped to protein-coding and long noncoding RNA genes. Analysis of the distribution of R-loop peaks at protein-coding and lncRNA genes between N2KO and Control cells showed that in both cell types R-loop signals were predominantly enriched at TSSs, with N2KO cells exhibiting significantly stronger enrichment at pre-existing loci and numerous gains of new TSS-associated R-loop signal (Fig. 4D). Importantly, this TSS-associated R-loop gain is not a byproduct of increased gene expression, as the genes harboring R-loop peaks (n = 225) were not found to be differentially expressed between N2KO and control cells by RNA-seq analysis (Fig. S4D). However, R-loop–associated genes are enriched in genes with high baseline expression (Fig. S4D) with housekeeping genes significantly enriched among the R-loop positive gene set (Odds Ratio=3.96, p=5.44×10^−6^, Fisher’s exact test). Overall, these data provide direct evidence that NEIL2 depletion results in TRC-related R-loops at TSS regions of actively transcribed genes.

Since loss of NEIL2 had a major impact on RNAPII dynamics in PDAC cells, we mapped the transcription impediment caused by NEIL2 deficiency by performing a ChIP-seq for elongating RNAPII CTD-S2 in Panc1-Cas9 Control and N2KO cells (Pol2S2 ChIP-seq). As expected from an elongation-associated modification, RNAPII CTD-S2 signal progressively accumulates towards gene ends (Fig 4F). Interestingly, a global comparison of Pol2S2 signals between Control and N2KO cells also revealed a marked increase of Pol2S2 signal within the ∼2 kb region downstream of the TES in N2KO cells (Fig. 4F). Analysis of the Pol2S2 signal in the 2kb region downstream of the TES identified 2974 genes exhibiting a >2-fold Pol2S2 accumulation in N2KO cells compared to Controls. Notably, the vast majority (∼94%) of genes exhibiting >2 kb of Pol2S2 accumulation downstream of the TES were not transcriptionally upregulated (Fig S4D), indicating that the increased Pol2S2 signal in N2KO is not simply a consequence of elevated transcriptional activity but instead reflects a distinct underlying mechanism. Specific examples of such Pol2S2 accumulation downstream the TES are shown for the EGR1 and CCN1 loci (Fig. 4E); in both cases, the Pol2S2 signal appears to reflect stalling of RNA Pol II during the termination phase of the respective genes in N2KO cells, rather than increased expression or confounding read-through from adjacent transcriptional units. To validate our sequencing results, we performed ChIP- and R-ChIP-qPCR on the representative gene EGR1 (Fig. S4E). Consistent with genome-wide analyses, EGR1 exhibited enriched R-loop occupancy at TSS site and elongating RNAPII occupancy downstream its TES sites, confirming the sequencing-based observations. Overall, these data indicate that NEIL2 loss impairs transcription termination.

Supporting the role of NEIL2 in the TES site, multiple transcription termination-associated factors were associated with NEIL2 in the co-IP mass spectrometry dataset. Furthermore, NEIL2 loss caused aberrant association of several RNA 3’UTR factors with RNAPII complex. For example, it reduced the association between elongating RNAPII CTD-S2 and the co-transcriptional splicing factor U2AF2(*54*), while increasing interaction of elongating RNAPII with the PAS factor CPSF1(*55*) and APA regulator CPSF5(*56*). Co-IP analyses revealed impaired CPSF1–CPSF5 interaction (Fig. 4G). Collectively, these data indicate that NEIL2 loss alter elongating RNAPII association with RNA 3’-processing factors, which potentially contribute to RNAPII retention at TESs and impaired transcription termination. To gain further insight into the possible roles of NEIL2 in transcription termination, we first analyzed polyA-selected data focusing on signal downstream of gene 3’ ends where RNAPII S2 aberrantly accumulates in N2KO cells. However, we did not detect any significant differences in the read-through transcription past the annotated gene ends between the Cas9 control and N2KO cells, as exemplified by EGR1 and CCN1 (Fig. 4E). Similarly, sequencing of total RNA after ribosomal RNA(rRNA)-depletion did not reveal increased signal downstream of TES (Fig. S4F).

We next investigated whether aberrant RNAPII occupancy at TES contributes to TRCs. Terminating RNAPII complex exhibits a unique phosphorylation of T4 residue on RNAPII-CTD(*57*). PLA analysis between terminating RNAPII (CTD-T4) and PCNA revealed increased signal in NEIL2 KO cells, indicating enhanced spatial proximity between stalled terminating RNAPII and replication machinery (Fig.4H). This finding suggests that accumulation of terminating RNAPII complexes downstream of TES contributes to TRC formation upon NEIL2 loss, potentially exacerbating transcription-coupled genotoxic stress from retained RNAPII at TES downstream (Fig. 4B).

### Inhibition of NEIL2 suppress survival and growth of pancreatic cancer cells

Given the pronounced TRC accumulation and subsequent genotoxic stress observed following NEIL2 knockout, we next investigated the functional impact of NEIL2 loss on pancreatic cancer cell fitness. Colony formation assays and real-time cell proliferation analysis (RTCA) demonstrated a significant reduction in Panc1 cell survival and proliferation following NEIL2 knockout (Fig. 5A and 5B). Re-expression of full-length and C-terminal truncated (CTD) NEIL2 partially rescued the proliferation defect, highlighting the functional contribution of NEIL2 to pancreatic cancer cell fitness (Fig S5A). To further evaluate the impact of NEIL2 loss on tumor growth, we performed 3D spheroid formation assays using Panc1-Cas9 Control and N2KO cells. At 72 hours post-seeding, N2KO spheroids were smaller and displayed irregular morphology compared to controls. Over the subsequent 72 hours, N2KO spheroids exhibited significantly reduced growth, indicating that NEIL2 loss impairs pancreatic cancer cell proliferation in both 2D and 3D culture models (Fig 5C).

**Figure 5:**
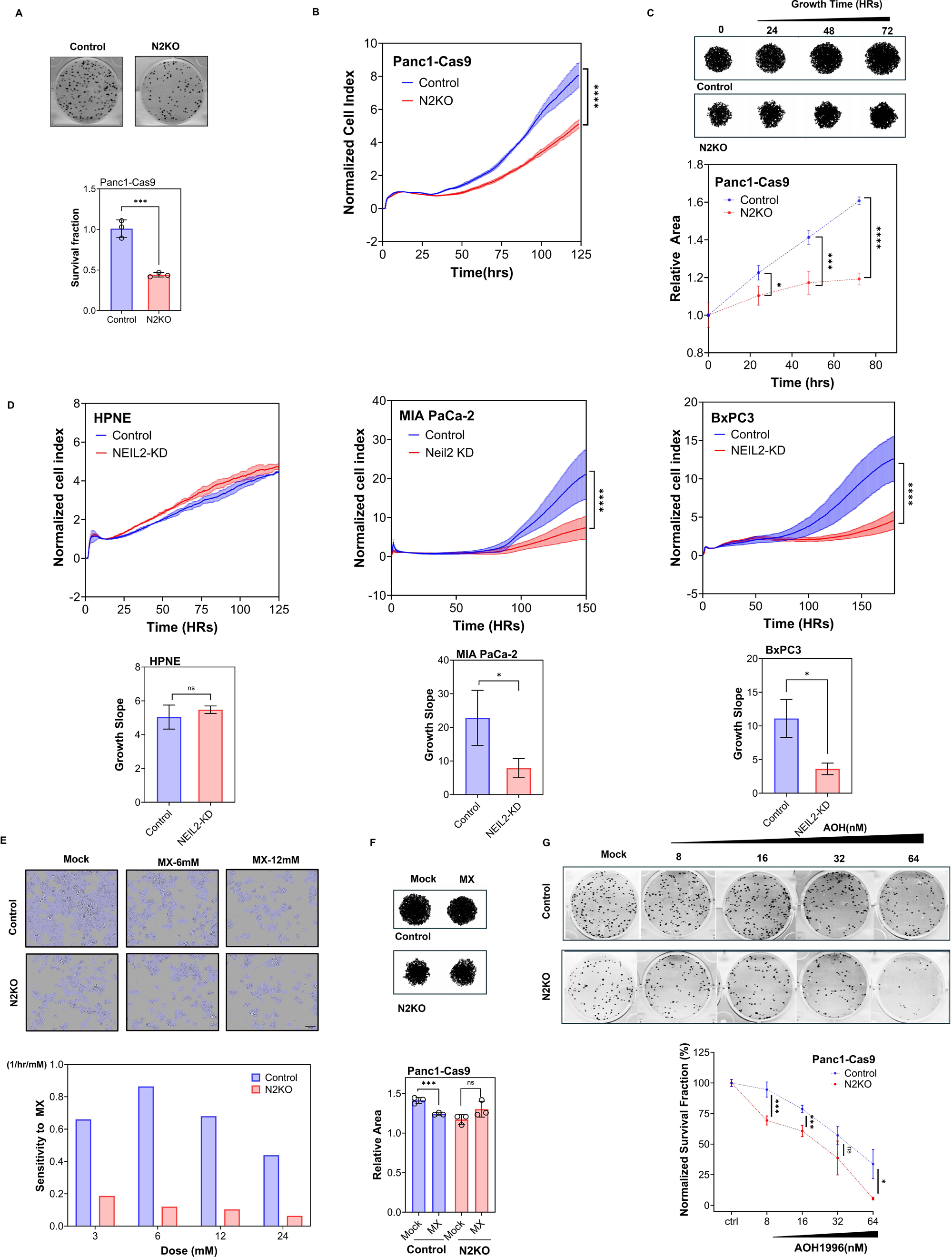
Inhibition of NEIL2 suppress survival and growth of pancreatic cancer cells. **(A)** Clonogenic survival assays comparing Panc1-Cas9 control(Control) and NEIL2-knockout(N2KO) cells. Representative colony formation images and quantification of survival fractions are shown. **(B)** Real-time proliferation analysis (RTCA) of Control and N2KO cells. Cell growth was monitored continuously; normalized cell index values are shown. **(C)** Three-dimensional spheroid growth assays in Control and N2KO cells. Representative binary spheroid images are shown at the indicated time points. Quantification of spheroid area was performed longitudinally to assess spheroid expansion and growth dynamics. Data are presented as mean spheroid area over time. **(D)** Growth kinetics following NEIL2 knockdown in HPNE, MiaPaCa2, and BxPC3 cells. Representative growth curves; calculated growth slopes are shown. **(E)** Image-based cell proliferation assays in Control and N2KO cells treated with increasing concentrations of methoxyamine (MX) as indicated. Cell growth was monitored by live-cell confluence, and MX sensitivity was quantified for each cell line using kinetic modeling. **(F)** Three-dimensional spheroid response to MX treatment. Representative spheroid images and quantification of spheroid area are shown. **(G)** Clonogenic survival following treatment with the PCNA inhibitor AOH1996. Representative colony formation assays and quantification are shown.

Notably, RTCA assays further revealed that growth inhibition caused by NEIL2 depletion, achieved via siRNA knockdown, was observed in PDAC cell lines, including MiaPaCa-2 and BxPC-3, but not in the non-transformed pancreatic epithelial cell line HPNE (Fig. 5D and Fig. S5B). Although driven by distinct oncogenic KRAS mutations, MIA PaCa-2 (KRAS G12C), PANC-1 (KRAS G12D), and BxPC-3 (KRAS WT) PDAC cells exhibit elevated basal TRC levels, whereas HPNE cells display a low TRC burden^11^. Consistently, PLA assays further confirmed that NEIL2 knockdown significantly increased TRCs in BxPC-3 and MIA PaCa-2 cells but caused no significant change in HPNE cells (Fig. S5C). These findings suggest that disrupting NEIL2-mediated TRC resolution may selectively impair pancreatic cancer cells while sparing normal pancreatic cells.

Since N2KO-associated TRCs were largely insensitive to BER inhibition by MX, we asked whether this saturation effect extends to cellular growth responses. Consistent with this hypothesis, real-time imaging assays showed that N2KO cells were less responsive to MX treatment across a range of concentrations. Kinetic modeling further indicated a significantly reduced affinity of N2KO cells for MX (Fig. 5E and Fig. S5D), Meanwhile, 3D spheroid assays showed that 72-hour MX treatment significantly reduced spheroid area in Panc1-Cas9 Control cells but had minimal effect on Panc1-Cas9 N2KO spheroids (Fig 5F). Together, our results support the conclusion that growth inhibition by MX in PDAC depends on intact NEIL2-mediated BER activity. The concordance between TRC levels and proliferation responses under MX treatment further reinforces the functional link between TRC regulation and pancreatic cancer cell growth.

We next explored whether exploiting N2KO-induced TRCs could enhance therapeutic vulnerability. To this end, we treated Panc1 cells with AOH1996, a PCNA-targeting compound that causes DNA damage in PDAC cells by stabilizing PCNA–RNAPII complexes at sites of TRCs. Colony formation assays revealed that N2KO cells were significantly sensitive to AOH1996 (*18, 58*), particularly at sublethal concentrations (8 nM) (Fig. 5G), accompanied by a moderate increase in late-apoptosis population (Fig. S5E). Collectively, these results demonstrate that disrupting NEIL2-mediated TRC resolution selectively suppresses pancreatic cancer cell proliferation and enhances sensitivity to TRC-targeting agent – AOH1996.

## Discussion

TRCs arise from disrupted spatial and temporal coordination between transcription and DNA replication machinery. Aberrant oncogenic signaling has been shown to promote TRCs and associated R-loop accumulation, leading to replication stress and transcription-coupled DNA damage. Examples include estrogen-induced replication stress in breast cancer and elevated TRCs driven by oncogenic HRAS^V12 (*59*). Our recent work showed that KRAS G12D induces a global increase in TRCs in pancreatic cancer cells, primarily through constitutive activation of ERK and AKT signaling (*7*). Although these studies have established strong mechanistic links between oncogenic signaling and TRC formation, most studies have relied on cell line–based models. Clinical relevance of TRCs and potential as therapeutic targets remains insufficiently defined.

In this study, we addressed this gap by applying a PLA-based TRC detection assay to patient-derived tissue microarrays. By analyzing paired tumors and adjacent normal tissues from untreated patients, we controlled for patient-specific measured and unmeasured confounders. Our results provide direct clinical evidence that pancreatic tumor tissues exhibit at least two-fold enrichment in TRCs compared to matched normal tissues. TRCs were also associated with features of aggressive disease, including larger tumor size and higher histologic grade. Although survival differences did not reach statistical significance, patients with high TRC burden showed shorter DFS and a tendency toward increased risk of disease recurrence and progression, suggesting a likely association between TRC accumulation and poor clinical outcomes. Using ICGC datasets, we previously demonstrated that promoter-proximal mutations in highly transcribed genes and common fragile site mutations, both associated with TRCs, are 30-to 120-fold more prevalent in PDAC than in breast, colon, and non-small cell lung cancers(*7*). These observations highlight TRCs as a common feature of aggressive PDAC biology and also as a potential therapeutic vulnerability.

TRCs represent an adaptive response to oncogene-driven replication and transcriptional dysregulation. Therefore, pathways that mitigate TRCs and R-loops likely play a central role in maintaining genomic integrity in tumor cells. At the same time, persistent TRCs represent an endogenous vulnerability that may be therapeutically exploited. We previously identified several BER factors, including NEIL2, POLB, APTX, FEN1, SOD1, PARP1, and APEX1, as contributors to TRC resolution. Although combined inhibition of the BER and ATR pathways showed synergistic therapeutic effects in PDAC models, significant toxicity of this combination could limit clinical application.

As TRC-targeting strategies continue to emerge, pharmacological agents that exploit TRC-associated vulnerabilities are gaining attention. The PCNA inhibitor AOH1996, which stabilizes PCNA–RNAPII interactions at sites of TRCs, has shown promising antitumor activity in PDAC and other solid tumors characterized by high TRC levels (*18*). These proof-of-concept studies have helped define a critical role of NEIL2 in TRC regulation in PDAC. Our data demonstrate that NEIL2 deficiency leads to accumulation of oxidative DNA lesions, including 8-oxo-dG. Although such lesions are traditionally considered non-helix distorting lesions and can be bypassed by RNAPII, studies propose that 8-oxo-dG in promoter regions can facilitate G-quadruplex formation (*60*), modulate transcription factor binding(*61*) and induce repressive histone modifications (*61, 62*), thereby regulating transcription initiation. Notably, several BER factors have emerged as direct regulators of transcription in cancer cells. For example, the DNA glycosylase OGG1 promotes estrogen-induced transcription by facilitating the removal of the repressive histone mark H3K9me2 (*63*). Similarly, the AP endonuclease APE1 directly binds G-quadruplex structures to enhance transcription of oncogenes, including KRAS, in pancreatic cancer cells (*64*). By rapidly sensing and resolving transcription-associated oxidative lesions and facilitating RNAPII initiation and termination, NEIL2 limits R-loop accumulation and suppresses TRC formation

Despite strong evidence supporting a BER-dependent role for NEIL2 in TRC regulation, our data also reveal noncanonical aspects of NEIL2 function. NEIL2 knockout attenuated the effects of BER inhibition on TRC accumulation and cell proliferation, indicating that NEIL2 operates upstream of BER-mediated TRC resolution. Surprisingly, a C-terminally truncated NEIL2 construct was sufficient to rescue TRC accumulation and proliferation defects, suggesting that the non-canonical function of NEIL2 extends beyond its glycosylase activity. Consistent with our findings, previous *in vitro* studies have shown that the NEIL2 C-terminal domain is essential for DNA lesion repair (*65–67*), but dispensable for protein–protein interactions (*48*). These results implicate the N-terminal protein–protein interaction domain of NEIL2 as a critical mediator of TRC and R-loop regulation.

In addition, our findings suggest that NEIL2 contributes to genome stability through the regulation of RNAPII dynamics during transcription termination. NEIL2 loss resulted in RNAPII accumulation near TESs without a corresponding increase in R-loop formation, indicating mechanisms distinct from the R-loop–dependent TRCs observed at TSSs. While previous studies have linked impaired topological stress resolution to transcriptional readthrough and replisome collisions (*68*), our data instead point to defects in 3′-end RNA processing, as NEIL2 deficiency disrupted the recruitment of alternative polyadenylation and co-transcriptional splicing factors, including CPSF1 and U2AF2. These findings suggest that NEIL2 may facilitate efficient transcription termination by coordinating RNA processing events required for timely RNAPII release from chromatin.

Although NEIL2 loss did not substantially alter global 3′ UTR usage or transcriptional readthrough, persistent RNAPII occupancy downstream of TESs may nevertheless increase opportunities for transcription-replication interference. Consistent with this possibility, we observed increased proximity between late-terminating RNAPII (CTD-T4) and PCNA in NEIL2-deficient cells, suggesting enhanced encounters between stalled transcription complexes and replication machinery. Together, these findings expand the role of NEIL2 beyond transcription-associated DNA repair and R-loop regulation, implicating NEIL2 in the maintenance of transcription termination fidelity. Whether termination-associated RNAPII stalling also contributes to TRC formation and genome instability remains an important question for future investigation.

More broadly, NEIL2 interactome places NEIL2 within a network of factors involved in transcription-coupled genome maintenance. Functional enrichment analyses of NEIL2-interacting TRC regulator revealed strong associations with RNA processing, DNA topology regulation, and DNA repair pathways, highlighting the close interplay between these processes during TRC resolution. Interestingly, our study provided preliminary evidence that the NEIL2-associated network may undergo cancer-specific rewiring, with several TRC regulators showing greater enrichment in PANC-1 cells than in normal pancreatic epithelial cells. This potential rewiring may reflect adaptation to increased transcriptional and replication stress and contribute to the heightened dependence of pancreatic cancer cells on NEIL2. Although NEIL2 knockout increased replication stress, slowed cell-cycle progression, and suppressed proliferation, it did not eliminate PDAC cells in our models. This suggests that compensatory DDR and anti-apoptotic pathways may be activated to buffer genotoxic stress resulting from NEIL2 deficiency. Comprehensive profiling of these compensatory DDR mechanisms will be critical to fully exploit NEIL2 as a therapeutic target.

In conclusion, our findings support a model in which NEIL2 mitigates TRC-mediated DNA damage in PDAC through its canonical TC-BER functions and DNA-binding independent regulation of RNAPII dynamics at TSS and TES. Our study identifies NEIL2 as a critical adaptive dependency and a therapeutic target that regulates TRCs in PDAC.

## Methods and Materials

### Cell line and Culture Condition

HPNE E6/E7/st, Panc-1, BxPC-3, Miapaca2, and HEK293T cells were obtained from commercially sourced (ATCC). Panc-1, Miapaca2, HEK293 were cultured with high-Glucose DMEM medium (Fisher, SH30423FS) supplemented with 10% FBS and 1% Pen-Strep (Sigma, 11074440001). The BxPC3 cells were cultured with RPMI1640 (Fisher, 11875119) supplemented with 10% FBS and 1% pen-strep. HPNE cells were cultured with filtered medium (75% DMEM low glucose, 25% M3 base medium, 5% fetal bovine serum (FBS), 10 ng/mL human recombinant EGF, 2 mM L-glutamine, 750 ng/mL puromycin).

### Plasmid Construction

To generate the NEIL2 C-terminally truncated mutant (NEIL2-CTDΔ) overexpressing vector, site-directed mutagenesis was performed on Flag-tagged full-length NEIL2 to truncate the NEIL2 C-terminal domain from Pro192 to Thr332 with primer pairs: 5’-gcgtacgcgttgtgaccaccggggaagag-3’ and 5’-ctcttccccggtggtcacaacgcgtacgc-3’. The NEIL2-CTDΔ construct was verified by sequencing, and protein expression was verified by immunoblotting. The NEIL2-CTDΔ construct and the full-length NEIL2 expression vector were transiently transfected into indicated cells for 72 hours before being subjected to assays.

### Transient transfection with siRNA and vectors

Cells were seeded at least 12 hours before transfection until they reached a confluency of 50% - 60%. For 6-well plates, 3.6 μL Lipofectamine 3000 was dissolved in 120 μL OPT-MEM, and 1:1 added to OPT-MEM medium containing 1.2 μg desired DNA vectors with 2.4 μL P3000 or Target-on NEIL2 siRNA at a final concentration of 50 nM without P3000. The transfection mix was then added to cells and incubated for 72 hours before being subjected to sequential assays. For cell proliferation assay, reverse transfection methods were applied; the lipofectamine-siRNA mix was pre-added to the cell suspension before seeding to plates.

### CRISPR-Cas9 genetic engineering

Panc1 cancer cells transfected with lentivirus prepackaged with inducible-Cas9 system were obtained from the Integrative Genomics Core at City of Hope. Cas9 expression was induced by 2 μg/mL doxycycline and confirmed by western blot. The established CRISPR-Cas9 Panc1 cell line was then transfected with multi-guided sgRNAs and cultured with 2 μg/mL doxycycline for 72 hours; sgRNA targeting sequences are listed in supplemental table 1. The NEIL2 sgRNAs transfected cell lines were then seeded into 96 wells at an average of 0.5 cells per well for limited dilution to select stable NEIL2 KO cell clones. NEIL2 KO efficiency was confirmed with immunoblotting and Sanger sequencing. Single NEIL2-deficient clones and NEIL2-deficient cell pools were generated to exclude the phenotype of interest due to clonal variation. In parallel, scrambled sgRNA was transfected to CRISPR-Cas9 cells as a negative control, indicated as Control. Both N2KO and Control cells were cultured in DMEM medium containing 10% Tet-free FBS.

### Western blot

A standard western blot protocol was used. Briefly, treated cells were lysed in non-SDS RIPA buffer (10 mM Tris-HCl pH 8.0, 1 mM EDTA, 0.5 mM EGTA, 1% Triton-X 100, 0.1% Sodium Deoxycholate, 140 mM NaCl) supplemented with protease and phosphatase inhibitor cocktail. After BCA assay quantification, cell lysates were mixed with equal volume of 2X Laemmli sampling buffer and loaded on 4–12% Tris-glycine gel for electrophoresis. The separated proteins were transferred onto PVDF membrane. The target bands were sequentially probed by primary antibodies and secondary antibody as indicated in the specific results with dilutions ranging from 1:1000 to 1:5000 (the antibody source and dilutions are indicated in the reagent/antibody list). The probed bands were revealed with West Pico plus luminol buffer (Fisher, 1863096).

### Reverse Transcription quantitative polymerase chain reaction

For RT–qPCR, total RNA was directly extracted and reverse transcribed by Cell-direct RT-qPCR kit. For ChIP-qPCR, the template DNA was obtained by the ethanol-precipitation step. RT–qPCR was performed using the PowerUp SYBR Green PCR Kit on Invitrogen PCR instrument with corresponding reaction conditions. The primers used are listed in the Supplementary Table 1.

### Co-immunoprecipitation (Co-IP) and mass-spectrometry (MS) analysis

To identify protein partners with NEIL2-WT and NEIL2-ΔCTD vectors, the Flag-tagged NEIL2 constructs, and empty pCMV-Flag negative control vectors were transfected into Panc1 cells and cultured for 72 hours. Cells were then lysed at 4°C for 30 mins with TBS buffer (50 mM Tris-HCl pH 7.5, 1 mM EDTA, 150 mM NaCl) supplemented with 1% Triton X-100 and protease inhibitor cocktail. The lysates were centrifuged at 14,000 rpm at 4°C for 10 mins, then incubated with Pierce M2 Flag affinity gel at 4°C overnight. The gel with pull-down beads was washed three times with TBS buffer supplemented with 0.5% Triton X-100 before being subjected to following assays.

For mass spectroscopy, washed M2 gel underwent on-gel digestion to extract pull-down proteins for sequential mass spectroscopy and data processing by Taplin mass spectrum facility. Briefly, M2 gel-bound proteins were washed extensively with 50 mM ammonium bicarbonate and digested on-bead overnight at 37°C using modified sequencing-grade trypsin (Promega). Peptides were recovered, dried in a SpeedVac, desalted using StageTips, and analyzed by nanoLC–MS/MS on a Velos Orbitrap Elite mass spectrometer (Thermo Fisher Scientific). MS/MS spectra were searched against the appropriate protein database using SEQUEST (Thermo Fisher Scientific), including a reversed decoy database for false discovery rate estimation. Peptide-spectrum matches were filtered to achieve a peptide false discovery rate of 1–2%. For quantitative analysis, to maximize the inclusion of potential components of the Neil2 interactomes, we selected proteins that were either uniquely detected in the NEIL2 Pulldown or enriched by at least two-fold relative to the negative control on summed intensity, with more than 1 unique peptide match with M/S ratio were counted. The relative protein abundance of two independent experiments was normalized by the Flag control IgG respectively, and plotted to characterize highly enriched NEIL2-associated proteins in both experiments. Proteins of interest were validated by Co-IP western blots. Gene ontology analyses were performed using enrichr(*69*) after filtering to retain proteins with an IP/control summed intensity ratio >10 and remove common contaminants (excluding proteins appearing in over 33% of control datasets in the CRAPome database(*70*)). Further functional enrichment analysis and network visualization of the NEIL2 interactome were performed using Metascape(*71*) based on the indicated filtered protein list.

### UNC cohort analysis

Bulk RNA-seq data from the UNC cohort were used to compare NEIL2 expression between primary PDAC tumors and PDAC metastases. NEIL2 expression was quantified as log₂-transformed transcripts per million (TPM) and visualized using violin plots.

PDAC single-cell RNA-seq data from the UNC cohort(*72*) were used to compare NEIL2 expression between the basal-like and classical transcriptomic subtypes. Tumor cells were reclustered using SCISSORS(*73*) with 3,000 highly variable genes, 20 principal components, a clustering resolution of 0.3, and a cosine distance metric. The number of nearest neighbors was set to the rounded square root of the total number of cells. Reclustered tumor cells were classified as basal-like or classical using PurIST(*74*), SCISSORS(*73*), and Elyada et al.(*75*) marker signatures. SCT-normalized NEIL2 expression was visualized using Seurat(*76*) (v5.3.0) and ggplot2(*77*) (v4.0.3). Differences between groups were assessed using two-sided Wilcoxon rank-sum tests.

### RNA-seq

PANC-1 Cas9 control and PANC1-N2KO cells were seeded in 6-well plates and cultured for 48 h to ∼80% confluence (∼1 × 10 cells per well). Three biological replicates were prepared for each cell line. Total RNA was isolated using TRIzol Reagent according to the manufacturer’s instructions with minor modifications. Briefly, cells were lysed in 1 mL TRIzol and incubated for 5 mins at room temperature, followed by addition of 200 μL chloroform. Samples were vigorously mixed, incubated for 2–3 min, and centrifuged at 12,000 × g for 15 mins at 4 °C to separate phases. The aqueous phase was transferred to a fresh tube, and RNA was precipitated with isopropanol at −80 °C for 1 h, followed by centrifugation at 12,000 × g for 10 min at 4 °C. The RNA pellet was washed with 70% ethanol, briefly air-dried, and resuspended in RNase-free water. RNA quality assessment, genomic DNA removal, library preparation, and sequencing were performed by Novogene (CA, USA). Downstream processing and analyses are described in the bioinformatics analysis section.

### ChIP/R-ChIP and data analysis

The ChIP experiments were performed based on a standard protocol. Briefly, 1 × 10 cells were fixed with 1% formaldehyde for 10–15 min at room temperature, then quenched by 125 mM Glycine for 15 min at room temperature. The fixed Panc1 cells were washed twice with DPBS, then subjected to cellular fractionation by incubation with cell lysis buffer (10 mM Tris-HCl pH 8.0, 10 mM NaCl, 0.5% NP-40 and 1× protease inhibitor cocktail) for 20 mins on ice, and then nuclei fraction was suspended in nuclei lysis buffer (50 mM Tris-HCl pH 8.0, 10 mM EDTA, 1% SDS and 1× protease inhibitor cocktail). Chromatin DNA was sheared to 200–600 bp in size by water-based sonication. Five percent chromatin fragment was saved as input and the remainder was incubated with Pierce A/G magnetic beads conjugated with anti-V5 antibody overnight at 4°C. The pull-down DNA-protein lysates were sequentially washed once with TSEI (20 mM Tris-HCl pH 8.0, 150 mM NaCl, 1% Triton X-100, 0.1% SDS, 2 mM EDTA and 1× protease inhibitor cocktail), once with TSEII (20 mM Tris-HCl pH 8.0, 500 mM NaCl, 1% Triton X-100, 0.1% SDS, 2 mM EDTA and 1× protease inhibitor cocktail), once with TSEIII (10 mM Tris-HCl pH 8.0, 250 mM LiCl, 1% NP-40, 1% Deoxycholate, 1 mM EDTA and 1× protease inhibitor cocktail) and once with TE buffer (10 mM Tris-HCl pH 8.0 and 1 mM EDTA). The protein-chromatin complex was eluted with elution buffer (10 mM Tris-HCl pH 8.0, 1% SDS and 1 mM EDTA) and decrosslinked by incubation overnight at 65°C. After sequential RNase A and Proteinase K treatment, the precipitated hybrid fragment was cleaned by phenol twice and phenol:chloroform:isoamyl alcohol once, followed by ethanol precipitation. The recovered fragment was either subjected to qPCR or library construction and sequencing conducted by Innomic (CA, USA). R-ChIP experiments were performed using the same ChIP protocol on Panc1 cells expressing RNASEH1/D210N to detect R-loops, and RNASEH1WKKD, which served as negative control.

### Bioinformatics Analysis of Omics Data

All next generation sequencing data was aligned to the human reference genome (hg38) and GENCODE v45 basic annotations using STAR retaining only uniquely mapping reads(*78*).Differential gene expression analysis was performed in R using the DESeq2 package(*79*). Peak calling was performed using MACS2 with default parameters(*80*). Signal intensities in specific regions were quantified using FeatureCounts(*81*), with ChIP signal calculated as the ratios of RPKM values in ChIP and input libraries. Plotting was performed using DeepTools(*82*), or customized R graphics.

### ICGC Somatic Mutation Analysis

Whole-genome somatic single-base substitution (SBS) calls and matched gene-expression data for PDAC were obtained from the International Cancer Genome Consortium (ICGC) Data Portal (*83*), specifically the Australian and Canadian pancreatic cancer projects (PACA-AU and PACA-CA; GRCh37) (*84*). Donors were retained if they had whole-genome SBS calls and matched *NEIL2* expression. For each donor, somatic substitutions in protein-coding genes were classified into the six pyrimidine-centered classes (C>A, C>G, C>T, T>A, T>C, T>G) and the fraction of each was computed over that donor’s total protein-coding-gene SBS count. *NEIL2* expression was taken as the per-donor normalized read count from ICGC RNA-sequencing data and averaged across specimens for donors with more than one. C>A was the pre-specified substitution class of interest, as C>A is the main mutation caused by unrepaired 8-oxoguanine (*85*). Per-donor C>A fraction was correlated with *NEIL2* expression by Spearman rank correlation. For visualization, donors were stratified by *NEIL2* expression quartiles computed within the analysis cohort (*NEIL2*-low, <=25th percentile; MID; *NEIL2*-high, >=75th percentile), and the *NEIL2*-low and *NEIL2*-high groups were compared by two-sided Mann-Whitney test. Analyses used Python with pandas (*86*) and scipy (*87*).

### PLA assays, flow cytometry, and immunocytochemistry

For high-throughput experiments, cells were seeded on collagen-coated 96-well plates. For validation experiments, cells were cultured on collagen-coated glass in 6-well plates. After reaching ∼60% to 70% confluence, cells were fixed with 4% paraformaldehyde (PFA) for 15 mins at room temperature. Fixed cells were washed twice with DPBS, then permeabilized by adding 0.5% NP-40 on ice for 5 mins. Subsequently, cells were washed with DPBS 3 times and blocked with 2% BSA-DPBS buffer at room temperature for 1 hour. Cells were then incubated with RNAPII CTD-S2 (1:500) and PCNA (1:500) antibodies at 4°C overnight. Proximity ligation was performed using the manufacturer’s instructions supplied with the kit (Sigma Aldrich). Samples were stained with DAPI, and images were acquired using ZEISS LSM900 confocal light microscopy. Images were processed and analyzed with QuPath and ImageJ (RRID:SCR_003070) to calculate the PLA mean intensity per nucleus in each captured image.

For staining tissue microarray (TMA), paraffin-embedded TMA slides were incubated in the following order: Xylene 10 mins twice, 100% ethanol 10 mins twice, 95% ethanol 10 mins once, 70% ethanol 10 mins once, 50% ethanol 10 mins once. Rehydrated slides were rinsed in ddH2O for 4 mins and washed with DPBS, followed by immersion in sodium citrate buffer (10 mM sodium citrate, 0.05% Tween 20, pH 6.0) for 45 min at 95 °C for antigen retrieval. Slides were next treated with 2% BSA-DPBS with 0.3% Triton X-100 for 1 hour at room temperature. Remaining steps were performed consistently with cultured cells on coverslips besides adjusting primary antibody dilution to 1:50–1:100 at incubation steps.

For flow cytometry PLA assay, treated cells at 60–70% confluence were trypsinized, washed at least once with DPBS, then fixed with 4% PFA for 15 mins followed by permeabilization with 5-minute incubation with 0.5% NP-40. Fixed cell pellets were collected after centrifugation at 400×g for 5 mins and washed with DPBS twice, blocked with commercial blocking buffer (supplied in kit) at room temperature for 1 hour, and then incubated with RNAPII-CTD S2 (1:100) and PCNA (1:100) antibodies overnight at 4°C. PLA reactions were performed using manufacturer instructions (Sigma Aldrich). PLA-fluorescent labeled cells were passed through a 100 µm strainer, and PLA signal was measured using Attune NXT cytometer. FlowJo (RRID:SCR_008520) was used for data processing, analysis, and visualization.

### Clonogenic assay

For clonogenic assays in 6-well plates, 400 cells were seeded per well with full medium, in triplicate wells per condition. After 2 hours of cell seeding, medium containing indicated drug dosage or mock condition was added dropwise. Cells were cultured with or without treatment for 12 days, followed by fixation and staining with 200 μL crystal violet-methanol buffer for 30 mins. After rinsing with tap water, colonies (cell number >50) from each well were counted and labeled under light microscopy. Plating efficiency (PE) and survival fraction (SF) were calculated by averaging control and treated cell colonies per well for each condition.

### DNA fiber assay

Panc1-Cas9 control and Panc1-N2KO cells were first subjected to sequential pulse labeling with 30 μM IdU for 1 hour. Then, cells were washed with fresh medium and incubated in medium containing 30 μM CIdU and indicated treatments for an additional 1 hour. After pulse labeling, cells were harvested and embedded in agarose plugs (TopVision, Thermo Fisher Scientific). These cells were lysed at 50°C in a solution containing 1% n-lauroylsarcosine, 0.5 M EDTA (pH 8.0), and 0.2 mg/mL proteinase K for 72 hours. Subsequently, lysed cells were washed with TE 8 buffer, treated with 200 μM PMSF, then washed again with TE 8 buffer before being stored in modified TE buffer [10 mM Tris (pH 8.0) and 50 mM EDTA]. Agarose plugs were melted, and agarose digested using β-Agarase (New England Biolabs). DNA was stretched on microscope slides coated with 3-aminopropyltriethoxysilane (Sigma–Aldrich). Stretched DNA was denatured in alkali buffer [0.1 M NaOH in 70% ethanol, 0.1% β-mercaptoethanol] and fixed with 0.5% glutaraldehyde. Slides were blocked with 1% bovine serum albumin for 20 min. IdU and CIdU were detected using mouse anti-IdU monoclonal antibody (BD) (1:100) and rat anti-CIdU monoclonal antibody (Abcam) (1:500), respectively, followed by Alexa Fluor 568-conjugated goat anti-mouse (Invitrogen) (1:100) and Dylight 488-conjugated goat anti-rat (1:50) antibodies. DNA fibers were examined using fluorescence microscopy (ZEISS Axio Observer).

### Flow cytometry apoptosis assay

Treated cells were cultured in 6-well plates and trypsinized at 70%–80% confluence. Cells were washed twice in cold phosphate-buffered saline (PBS) and counted. Resuspended cells were adjusted to ∼300,000 cells per sample, stained with 100 μL propidium iodide (PI) in HBSS (ThermoFisher, 14175095) at 1:500 dilution, then 5 μL Annexin V-FITC was added for 15 minutes incubation at room temperature. After incubation, 400 μL HBSS was added to each sample and mixed gently. Apoptotic activity was measured within 15 minutes after staining using NovoCyte Quanteon Flow Cytometer (Agilent, USA). 50,000 events were quantified per sample. Results were presented as mean ± standard error in technical triplicates.

### Cell proliferation with real-time cell analysis (RTCA) and live-cell-image analysis

To measure the inhibitory effect of methoxyamine (MX) and ATR inhibitor (VX970) in Panc1 and MIA PaCa2 cells, RTCA assay was used to measure real-time cell growth. Approximately 4,000 cells were seeded on an E-plate (Agilent 05469830001). Cell growth curves were recorded on xCELLigence RTCA DP (Agilent, CA, USA) at 15-min intervals under standard cell culture conditions (37°C, 5% CO2). Indicated treatments were added 24 hours after plating. Cell index reflects cell numbers, and cell growth slope was calculated by the built-in algorithm in GraphPad prism.

### 3D spheroid formation assay

PANC-1 cells were cultured to 70–80% confluency and resuspended at 2 × 10 cells/mL. Cells were seeded into low-attachment U-bottom 96-well plates (Thermo Fisher Scientific, Cat# 174925) by adding 20 µL cell suspension to 130 µL spheroid medium per well (4,000 cells/well; 150 µL total volume). Spheroid medium consisted of DMEM supplemented with 10% FBS and EGF (40 ng/mL). Plates were centrifuged at 300 × g for 5 min and incubated at 37°C, 5% CO₂ for 72 h to allow spheroid formation. Sequential treatment conditions (dosage and duration) were applied as indicated in the corresponding figure. Spheroids were imaged every 24 h, and area was quantified using ImageJ (NIH).

### Quantification and statistical analysis

For TRC-PLA and immunofluorescence data, flow cytometry data, DNA fiber analysis, and colony formation assay (statistical analyses were performed using Prism 10.0 (GraphPad Prism, RRID:SCR_002798). Data are presented as mean ± SD, unless otherwise indicated. Differences between two groups were evaluated using unpaired Student’s t-test. P < 0.05 was defined as statistically significant. For RTCA data, Tukey’s test was used for multiple comparisons after one-way ANOVA. Formal power analysis was not performed. Data are representative of at least three independent biological experiments. P < 0.05 (*), P < 0.01 (**), P < 0.001 (***), P < 0.0001 (****); ns, not significant.

### Kinetic modeling

To quantify sensitivity of N2KO and Control cells to MX, we fit a kinetic model of the form 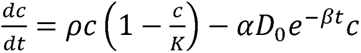 to untreated and treated in vitro data. The model combines logistic growth of the cells with a mass action killing effect of MX, with cell growth rate *ρ* (1/hour), MX dose *D*_0_ (mM) and decay constant 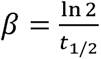 with half-life *t*_1/2_ = 40.2 (hours), and MX sensitivity parameter *α* (1/hour/mM). The cell growth rate constant *ρ* was estimated by fitting the model to untreated data for each line with *D*_0_ = 0, yielding *ρ_Cas_*_9_ = 0.038 (1/hour) and *ρ_N2KO_* = 0.021 (1/hour). The administered dose *D*_0_ was determined experimentally (*D*_0_ ∈ {3, 6, 12, 24} *mM*), and the cell growth *ρ* and MX clearance rate *β* were held constant for all conditions to yield a value of the sensitivity parameter *α* for each cell line. All parameter estimations used constrained nonlinear multivariable minimization (fmincon) in MATLAB version 26.1.0.3251617 (R2026a Update 2). MX sensitivity values were scaled by the cell line growth rate for comparison, revealing an increased sensitivity Control cells to MX as compared to N2KO (Figure 5C).

## Data availability

Requests for materials should be addressed to the corresponding authors. The mass spectrometry proteomics data have been deposited to the ProteomeXchange via the PRIDE repository with the dataset identifier PXD085239. RNA-seq, ChIP-seq, and R-ChIP-seq data have been deposited in the NCBI Gene Expression Omnibus (GEO) under accession numbers GSE350338 (poly(A)-selected RNA-seq), GSE350339 (total RNA-seq), GSE350337 (ChIP-seq), and GSE350322 (R-ChIP-seq). All other data generated in this study are provided within the Article and its Supplementary Information. Source data are provided with this paper.

## Author Contribution

F.M., M.N. and M.R. conceived and designed the study. F.M. performed the majority of the cellular and molecular biology experiments. M.N. performed the bioinformatic analysis of the sequencing data. F.M. and M.N. conducted the proteomic analysis. F.M., M.N. and M.R. analyzed the data and prepared the figures. A.F. and T.Z. performed and analyzed the DNA fiber assays. C.L. and J.J.Y. performed the UNC cohort analysis. M.A. performed the ICGC somatic mutation analysis. Q.F.performed the apoptosis assays and contributed to data analysis. A.L. and R.R. conducted kinetic modeling. H.F. assisted with molecular cloning and gene editing. T.L. assisted with cellular experiments and data analysis. D.D. provided clinical samples, pathological evaluation and clinical data. F.M., M.N. and M.R. wrote, revised and finalized the manuscript, with writing assistance from Y,L. T.W., B.S., Y.R.L., and Y.L. provided intellectual input, resources, and supervision of their respective contributions to the study. M.R. supervised the study, coordinated the collaborations and acquired funding.

## Competing Interests Declarations

The authors declare no competing interests

## Ethics Declarations

The construction and use of TMA was conducted in accordance with the U.S. Common Rule and the ethical principles of the Belmont Report. The construction and use of TMAs and patient records was performed under UCLA institutional review board approved protocol (IRB-11-0512) with an exemption for patient informed consent approved on the basis of the use of remnant pathology material and deidentified patient data (exempt category 4). The protocol was initially approved on February 14, 2011, and has been reviewed and approved annually, with current approval through February 8, 2027.

## Acknowledgement

We are grateful to Dr. Tapas K. Hazra (*The University of Texas Medical Branch, Galveston, TX, USA*) for providing the NEIL2 antibody that supported the initial phases of this work.

We are grateful to Dr. Linda Malkas (*City of Hope National Medical Center, Duarte, CA, USA*) for providing the AOH1996 compound for our functional study

We are grateful to Andrea Lynch (Beckman Research Institute of City of Hope, Duarte, CA, USA) for her assistance with reference management and citation formation.

This work was supported by the following funding sources awarded to M.R.: the Pancreatic Cancer Action Network Career Development Award (grant #20-20-RAOO), the National Comprehensive Cancer Network Young Investigator Award (grant #P062683), the American Cancer Society Institutional Research Grant (grant #132966-IRG), the City of Hope Academic Innovation Fund, and anonymous philanthropic support (GK & KK).

Research reported in this publication included work performed in the Integrative Genomics Core, Analytical Cytometry Shared Resource, Light Microscopy & Digital Imaging Shared Resource, which are supported by the National Cancer Institute of the National Institutes of Health under award number P30CA033572.

## Notes

### Competing Interest Statement

The authors have declared no competing interest.

