## supplementary figures and Captions for "NEIL2 regulates RNA polymerase II dynamics to suppress transcription–replication conflicts in pancreatic cancer"

**Supplementary Figure S1: Human PDAC shows enrichment of TRCs**

**(A)** Quantification of RNAPII–PCNA proximity ligation assay (PLA) foci in CK19-positive malignant epithelial cells, CK19-negative stromal cells, and adjacent normal pancreatic tissues from patient#112. **(B)** Correlation analysis between tumor size(cm) and relative PLA enrichment in tumor tissues. Linear regression analysis indicates a positive association between TRC accumulation and tumor size. Each dot represents an individual patient sample **(C)** Distribution of PLA-positive cells among PDAC specimens stratified by histological grade and Disease-free survival(DFS) of patients with PDAC stratified by tumor PLA burden. Kaplan–Meier analysis comparing the PLA-low and PLA-high groups using 25% PLA positivity as the cutoff. Differences were evaluated using the log-rank test. The number of patients at risk in each group at the indicated time points is shown below the Kaplan–Meier curve.**(D)** Tissue- and cell type-specific expression analysis of NEIL2 using transcriptomic data from the Human Protein Atlas. Dot plots depict NEIL2 expression across major human tissues and cell populations. Dot size reflects normalized transcript abundance of NEIL2. **(E)** NEIL2 expression across PDAC transcriptomic subtypes. Single-cell RNA-seq data were visualized to show basal-like and classical cell populations and NEIL2 expression. Violin plots showed no significant difference in NEIL2 expression between the two subtypes. ns, not significant; Wilcoxon rank-sum test.

**Supplementary Figure S2: NEIL2 deprivation leads to TRC accumulation and DNA damage in PDAC cells**

**(A)** NEIL2 interactome are ranked according to $\log_2$-transformed normalized total peptide counts. Selected TRC-associated factors are highlighted. The accompanying table lists representative NEIL2-interacting TRC regulators, including raw peptide counts and normalized abundance values. **(B)** Gene Ontology (GO) biological process enrichment analysis of proteins identified in both N2 and N2B immunoprecipitants. Proteins enriched relative to IgG controls (sum intensity/IgG > 10, CRAPome >33%) were subjected to GO analysis. Dot plots display significantly enriched biological pathways, including RNA metabolism, ribonucleoprotein complex biogenesis, DNA repair, transcriptional regulation, and genome maintenance. Dot size corresponds to the number of proteins associated with each term, and color intensity reflects enrichment significance. **(C)** Comparative proteomic analysis of the NEIL2 interactome in Panc1 and HPNE cells. The Venn diagram shows shared and cell type-specific NEIL2-associated proteins. The heatmap depicts the Panc1/HPNE ratio of normalized total peptide counts for known TRC regulators identified in the NEIL2 interactome. Differential abundance analysis is shown as a scatter plot with $log_2$-transformed normalized total peptide counts [$log_2$(normalized count + 1)] in HPNE and Panc1 cells plotted on the x- and y-axes, respectively. Preferential enrichment of TRC-associated proteins in Panc1 cells indicates cancer-specific remodeling of the NEIL2 interaction network.

**Supplementary Figure S3: NEIL2 resolves TRCs via TC-BER pathway but independent of DNA binding**

**(A)** Flow cytometric quantification of 8-oxo-dG levels during different cell-cycle phases in Panc1-Cas9 control(Control) and NEIL2-knockout(N2KO) cells. Representative FlowJo cell-cycle modeling plots and corresponding flow cytometry histograms are shown. G1, blue; S, yellow; G2/M, green. N2KO cells exhibited increased oxidative DNA damage throughout the cell cycle. **(B)** *NEIL2* expression inversely correlates with the C>A transversion fraction in pancreatic cancer. Per-tumor C>A fraction (of protein-coding-gene SBS) by *NEIL2* expression quartile in ICGC PDAC (n = 254: *NEIL2*-low $n$ = 64, MID $n$ = 126, *NEIL2*-high $n$ = 64). Boxes show median and IQR, points are individual donors. *NEIL2* correlates negatively with C>A fraction (Spearman $r = -0.18$, $p = 0.004$; *NEIL2*-low vs -high two-sided Mann-Whitney $p = 0.036$), consistent with the 8-oxo-dG accumulation in *NEIL2*-knockout cells (Fig. S3A). **(C)** DNA fiber assay analysis following treatment with N-acetylcysteine amide (NACA), aphidicolin (APH), or triptolide (TRP). Representative fiber images and quantification of replication track lengths demonstrate differential regulation of replication dynamics by modulating oxidative stress, transcription initiation, and replication stress. **(D)** Venn diagram illustrating the overlap of proteins identified by mass spectrometry in wild-type NEIL2 and the NEIL2 C-terminal deletion mutant (ΔCTD). Functional enrichment analysis identifies that 116 shared proteins associated with RNA processing, transcription regulation, and genome stability. **(E)** Co-immunoprecipitation analysis of exogenously expressed NEIL2 complexes. Immunoblotting confirmed interactions between wild-type NEIL2 and the NEIL2 C-terminal deletion mutant (ΔCTD) with RNAPII complex RBP1 CTD,

**Supplementary Figure S4: RNAPII dysregulation at TSS and TES drives TRCs in NEIL2-knockdown cells**

**(A)** Time-course PLA analysis of TRC resolution following transient aphidicolin treatment and recover. Representative PLA images and quantification reveal prolonged persistence of TRCs in NEIL2-knockout (N2KO) comparing with Panc1-cas9 control(Control) cells. Scale bar, 30 µm. **(B)** Analysis of TRC dynamics following transcription inhibition with triptolide and subsequent recovery. N2KO cells exhibit impaired resolution of TRCs during recovery. Representative images show PLA foci (red) and DAPI-stained nuclei (blue). Violin plots represent distributions of single-nucleus PLA intensities. Scale bar, 30 µm. **(C)** Dot blot analysis of genomic DNA–RNA hybrids isolated from Control and N2KO cells.

Indicated amounts of nuclear genomic DNA were spotted onto membranes and probed with the S9.6 antibody to detect DNA–RNA hybrids. Samples treated with RNase H prior to blotting served as specificity controls for hybrid detection. Representative dot blots and quantification demonstrate increased accumulation of DNA–RNA hybrids in N2KO cells relative to Control, consistent with elevated R-loop formation following NEIL2 loss. **(D)** Differential gene expression analysis in Control and N2KO cells. Violin plots compare baseline transcript abundance between genes exhibiting high and low R-loop occupancy, demonstrating the relationship between transcriptional activity and R-loop formation. The MA plot depicts transcriptome-wide expression changes following NEIL2 depletion, with significantly differentially expressed genes highlighted in green. Genes exhibiting increased elongating RNA polymerase II occupancy at transcription end sites (TESs) are highlighted in orange**. (E)** R-ChIP–qPCR and RNA polymerase II ChIP–qPCR validation at representative loci (EGR1 and CCN1). R-loop enrichment and elongating RNA polymerase II (POLR2S2) occupancy were quantified at transcription start sites (TSSs) and downstream regions of transcription end sites (TESs) by qPCR**. (F)** Scatter plot showing the correlation between changes in PolIIS2 enrichment and RNA-seq signal (rRNA-depleted libraries) 2kb downstream annotated TES of protein-coding genes. Data points are colored according to changes in the gene expression between N2KO and Panc1 control cells.

**Supplementary Figure S5: Inhibition of NEIL2 suppress survival and growth of pancreatic cancer cells**

**(A)** Real-time proliferation analysis of Panc1-Cas9 control (Control)cells transfected with FLAG control and Panc1-N2KO (N2KO) cells transfected with FLAG control, wild-type NEIL2, or the NEIL2 ΔCTD construct. **(B)** Validation of NEIL2 knockdown(KD) in HPNE, MiaPaCa2, and BxPC3 cells using independent siRNAs. Quantitative RT–PCR and immunoblot analyses confirm efficient depletion of NEIL2 expression. **(C)** PLA analysis of TRCs following NEIL2 KD in HPNE, MIA PaCa-2, and BXPC3 cells. Mock indicates cells transfected with a negative-control siRNA, and siN2 indicates cells transfected with a NEIL2-targeting siRNA. Scale bar, 30 μm. **(D)** Real-time image-based proliferation measurements of Control and N2KO cells treated with increasing concentrations of MX. Growth curves and calculated growth slopes demonstrate less sensitivity of N2KO cells. **(E)** Annexin V/propidium iodide flow cytometry analysis following treatment with the PCNA inhibitor AOH1996. Representative flow cytometry plots and quantification of viable, early apoptotic, late apoptotic, and necrotic cell populations are shown. N2KO cells exhibit modest enhanced apoptotic responses to PCNA inhibition.

**Fig.S1**

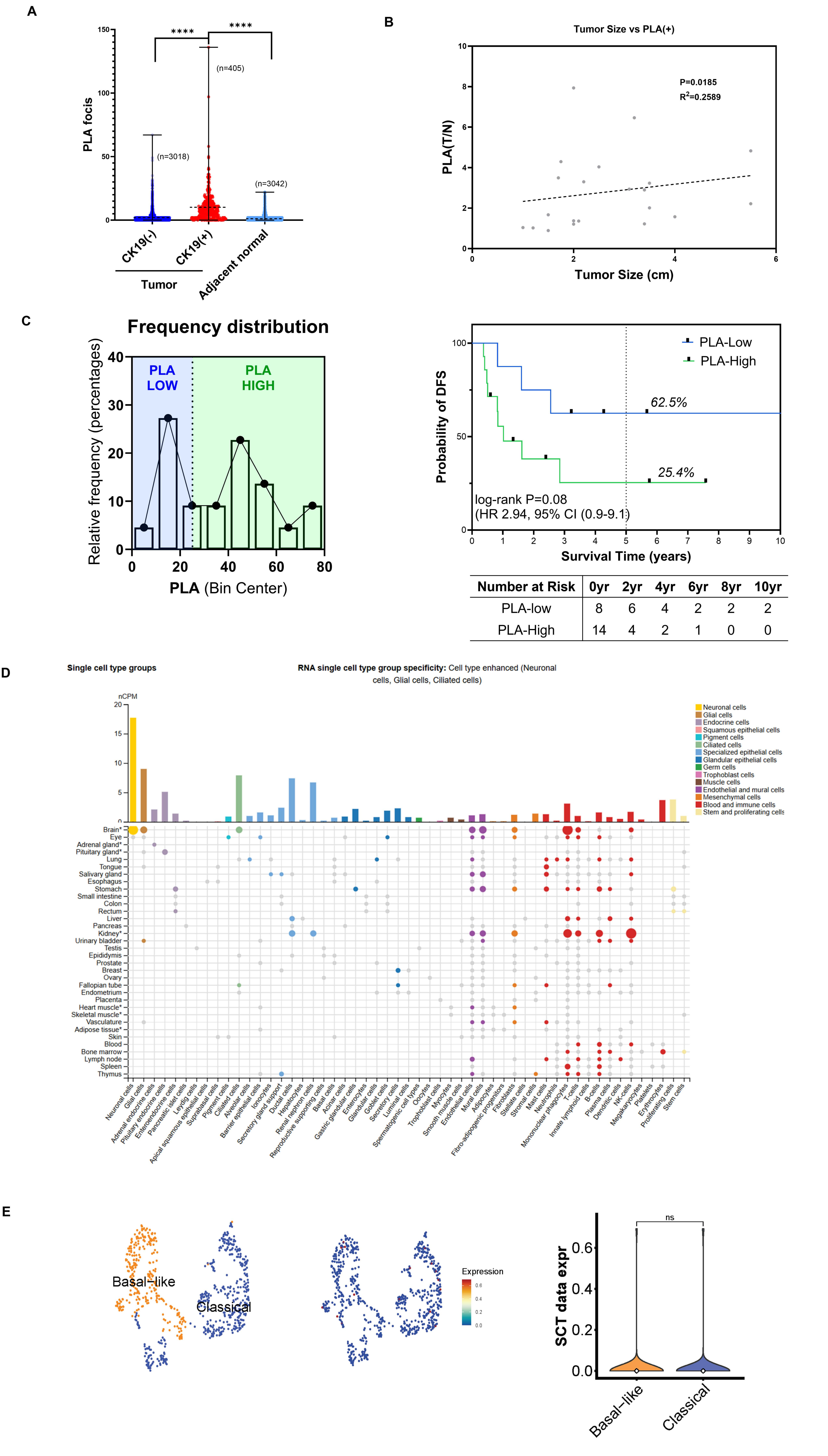

A
PLA focis
****
****
(n=3018)
(n=405)
(n=3042)
CK19(-)
CK19(+)
Tumor
Adjacent normal
B
Tumor Size vs PLA(+)
P=0.0185
R2=0.2589
PLA(T/N)
Tumor Size (cm)
C
Frequency distribution
PLA LOW
PLA HIGH
Relative frequency (percentages)
PLA (Bin Center)
PLA-Low
PLA-High
62.5%
25.4%
log-rank P=0.08
(HR 2.94, 95% CI (0.9-9.1)
Probability of DFS
Survival Time (years)
Number at Risk 0yr 2yr 4yr 6yr 8yr 10yr
PLA-low 8 6 4 2 2 2
PLA-High 14 4 2 1 0 0
D
Single cell type groups
RNA single cell type group specificity: Cell type enhanced (Neuronal cells, Glial cells, Ciliated cells)
nCPM
Neuronal cells
Glial cells
Endocrine cells
Squamous epithelial cells
Pigment cells
Ciliated cells
Specialized epithelial cells
Glandular epithelial cells
Germ cells
Trophoblast cells
Muscle cells
Endothelial and mural cells
Mesenchymal cells
Blood and immune cells
Stem and proliferating cells
E
Basal-like
Classical
Expression
ns
SCT data expr

**Fig.S2**

**A**

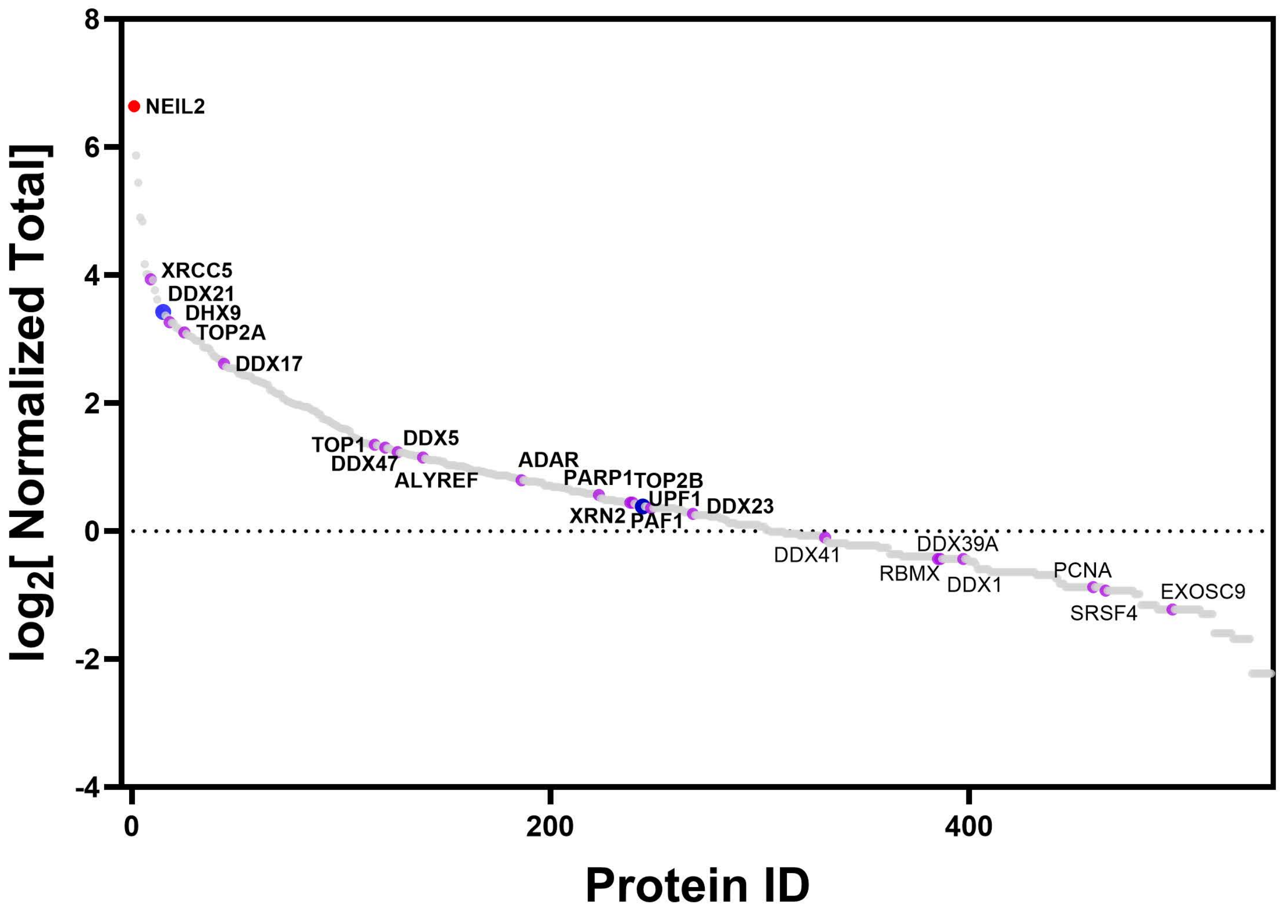


| Protein | Raw Total Counts | | | | Normalized | | |
|---|---|---|---|---|---|---|---|
| | Ctrl | N2 | N2B | Mean | N2 | N2B | Mean |
| NEIL2 | 0 | 426 | 515 | 470.5 | 100.0 | 100.0 | 100.0 |
| XRCC5 | 8 | 80 | 61 | 70.5 | 18.8 | 11.8 | 15.3 |
| DDX21 | 10 | 51 | 49 | 50 | 12.0 | 9.5 | 10.7 |
| DHX9 | 9 | 39 | 52 | 45.5 | 9.2 | 10.1 | 9.6 |
| TOP2A | 4 | 32 | 50 | 41 | 7.5 | 9.7 | 8.6 |
| DDX17 | 6 | 35 | 21 | 28 | 8.2 | 4.1 | 6.1 |
| TOP1 | 5 | 11 | 13 | 12 | 2.6 | 2.5 | 2.6 |
| DDX47 | 0 | 12 | 11 | 11.5 | 2.8 | 2.1 | 2.5 |
| DDX5 | 3 | 11 | 11 | 11 | 2.6 | 2.1 | 2.4 |
| ALYREF | 2 | 9 | 12 | 10.5 | 2.1 | 2.3 | 2.2 |
| ADAR | 4 | 9 | 7 | 8 | 2.1 | 1.4 | 1.7 |
| PARP1 | 1 | 6 | 8 | 7 | 1.4 | 1.6 | 1.5 |
| TOP2B | 0 | 5 | 8 | 6.5 | 1.2 | 1.6 | 1.4 |
| XRN2 | 1 | 5 | 8 | 6.5 | 1.2 | 1.6 | 1.4 |
| UPF1 | 3 | 7 | 5 | 6 | 1.6 | 1.0 | 0.7 |
| PAF1 | 2 | 6 | 6 | 6 | 1.4 | 1.2 | 1.3 |
| DDX23 | 0 | 7 | 4 | 5.5 | 1.6 | 0.8 | 1.2 |
| DDX41 | 0 | 3 | 6 | 4.5 | 0.7 | 1.2 | 0.9 |
| DDX1 | 0 | 3 | 4 | 3.5 | 0.7 | 0.8 | 0.7 |
| DDX39A | 1 | 3 | 4 | 3.5 | 0.7 | 0.8 | 0.7 |
| RBMX | 1 | 3 | 4 | 3.5 | 0.7 | 0.8 | 0.7 |
| PCNA | 1 | 2 | 3 | 2.5 | 0.5 | 0.6 | 0.5 |
| SRSF4 | 0 | 3 | 2 | 2.5 | 0.7 | 0.4 | 0.5 |
| EXOSC9 | 0 | 2 | 2 | 2 | 0.5 | 0.4 | 0.4 |

**B**

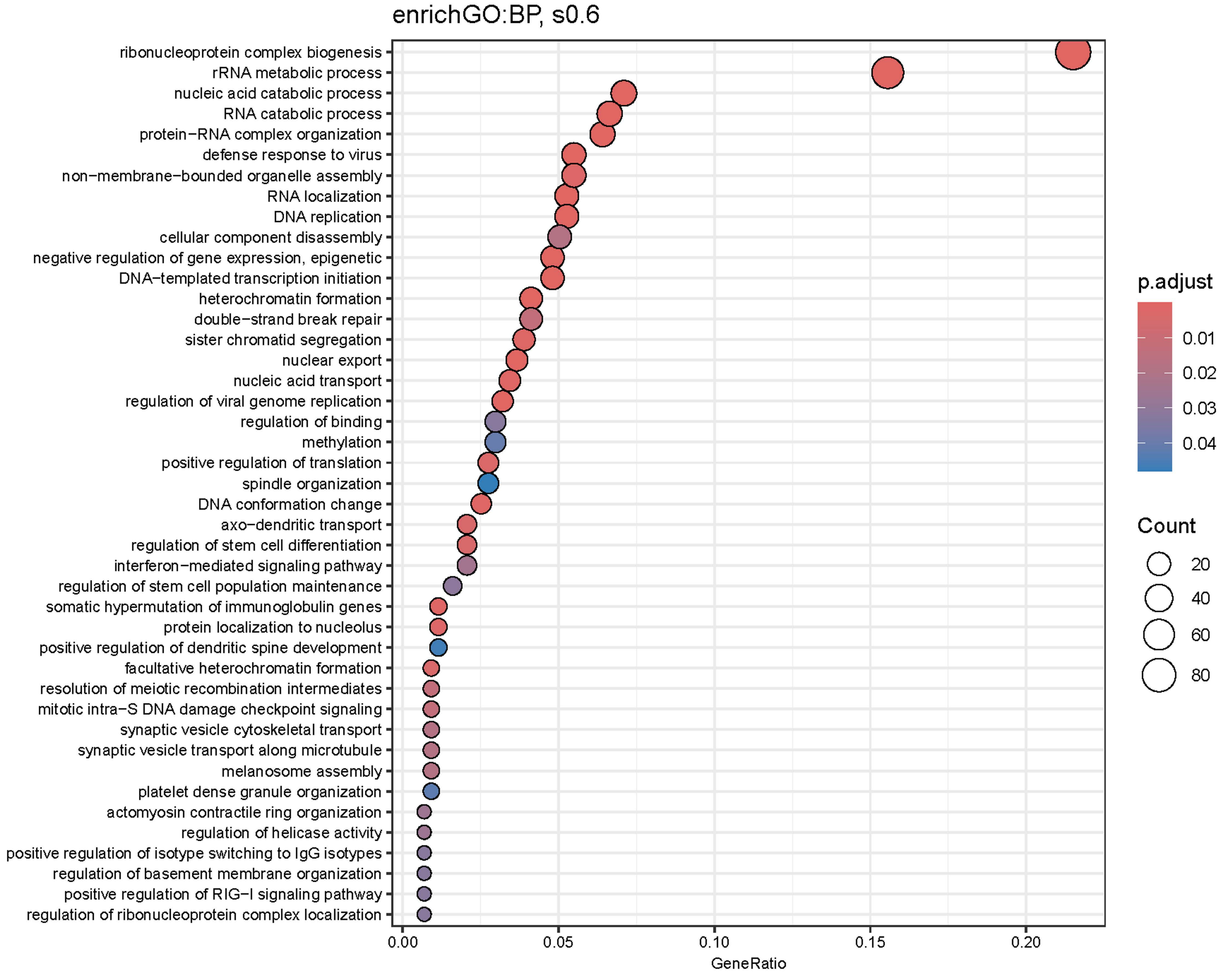


**C**

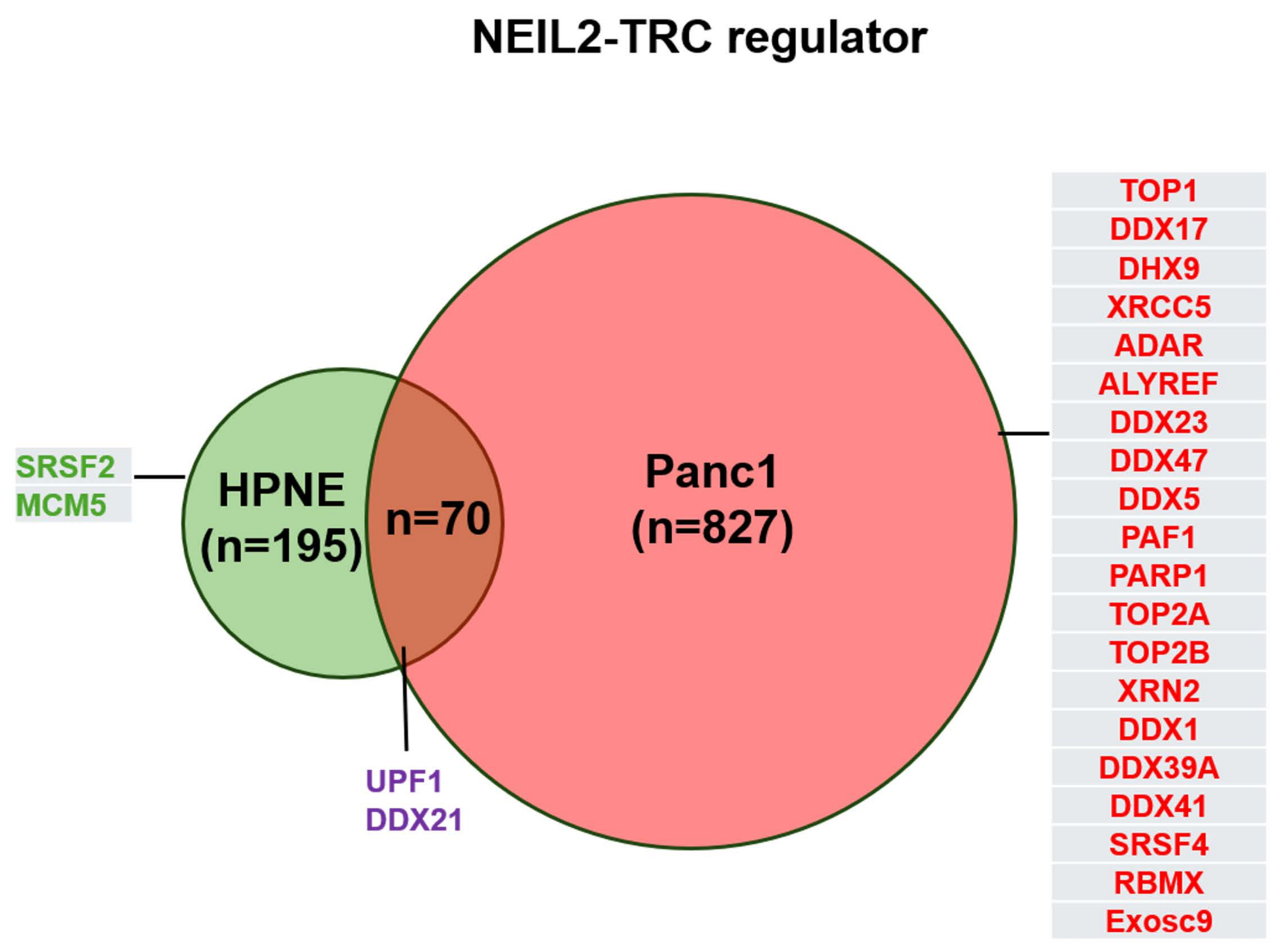


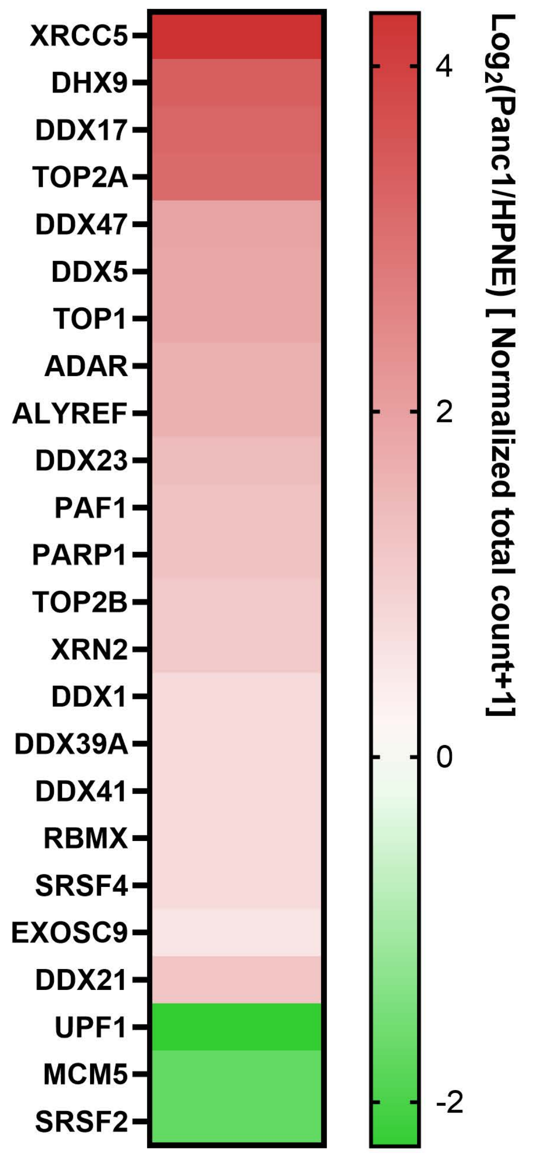


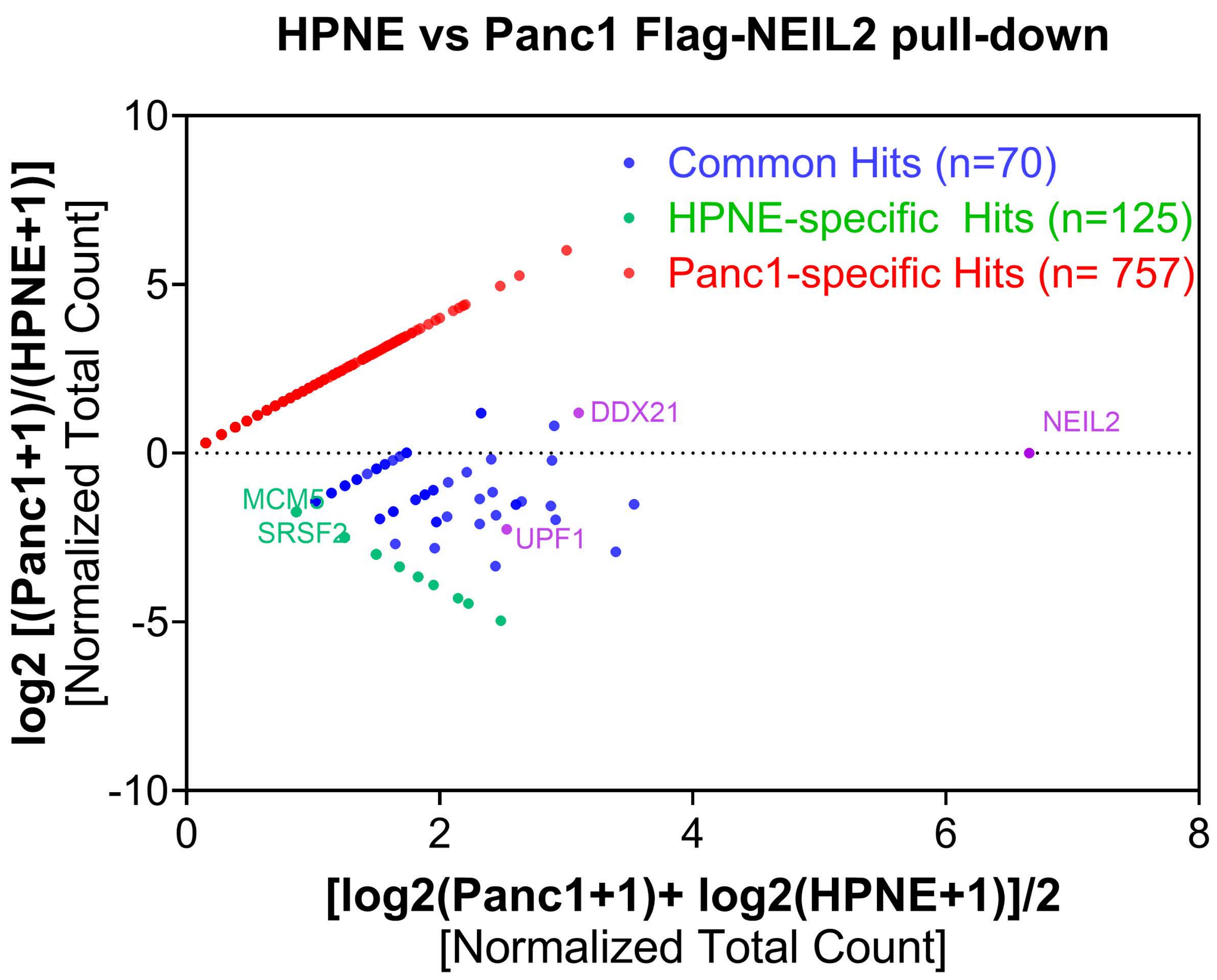

**Fig.S3**

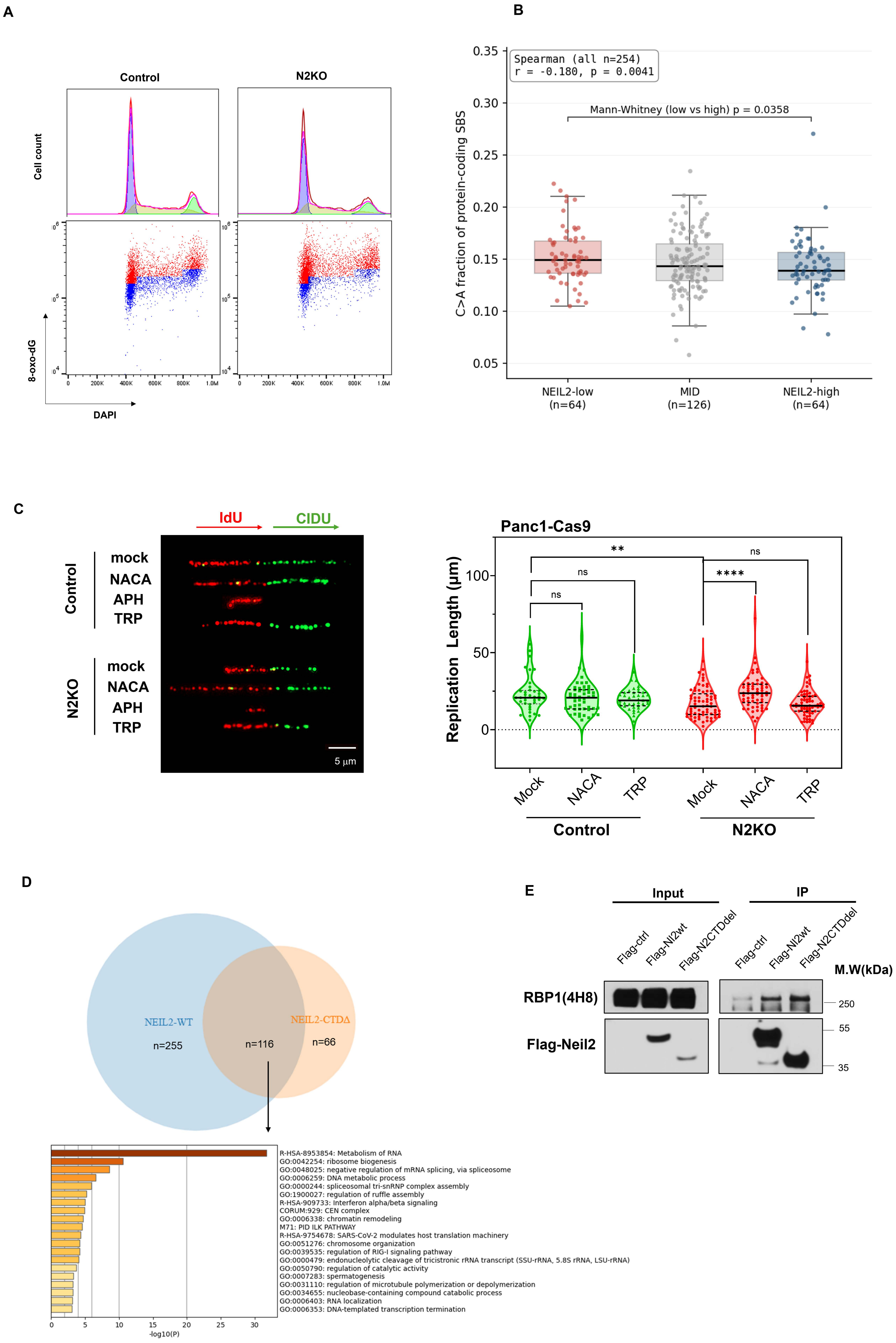
A
Control
N2KO
Cell count
8-oxo-dG
DAPI
B
Spearman (all n=254)
r = -0.180, p = 0.0041
Mann-Whitney (low vs high) p = 0.0358
C>A fraction of protein-coding SBS
NEIL2-low
(n=64)
MID
(n=126)
NEIL2-high
(n=64)
C
IdU
CIDU
Control
N2KO
mock
NACA
APH
TRP
5 μm
Panc1-Cas9
Replication Length (μm)
**
ns
****
Mock
NACA
TRP
D
NEIL2-WT
n=255
n=116
NEIL2-CTDΔ
n=66
R-HSA-8953854: Metabolism of RNA
GO:0042254: ribosome biogenesis
GO:0048025: negative regulation of mRNA splicing, via spliceosome
GO:0006259: DNA metabolic process
GO:0000244: spliceosomal tri-snRNP complex assembly
GO:1900027: regulation of ruffle assembly
R-HSA-909733: Interferon alpha/beta signaling
CORUM:929: CEN complex
GO:0006338: chromatin remodeling
M71: PID ILK PATHWAY
R-HSA-9754678: SARS-CoV-2 modulates host translation machinery
GO:0051276: chromosome organization
GO:0039535: regulation of RIG-I signaling pathway
GO:0000479: endonucleolytic cleavage of tricistronic rRNA transcript (SSU-rRNA, 5.8S rRNA, LSU-rRNA)
GO:0050790: regulation of catalytic activity
GO:0007283: spermatogenesis
GO:0031110: regulation of microtubule polymerization or depolymerization
GO:0034655: nucleobase-containing compound catabolic process
GO:0006403: RNA localization
GO:0006353: DNA-templated transcription termination
-log10(P)
E
Input
IP
Flag-ctrl
Flag-Nl2wt
Flag-N2CTDdel
M.W(kDa)
RBP1(4H8)
Flag-Neil2
250
55
35

Fig.S4

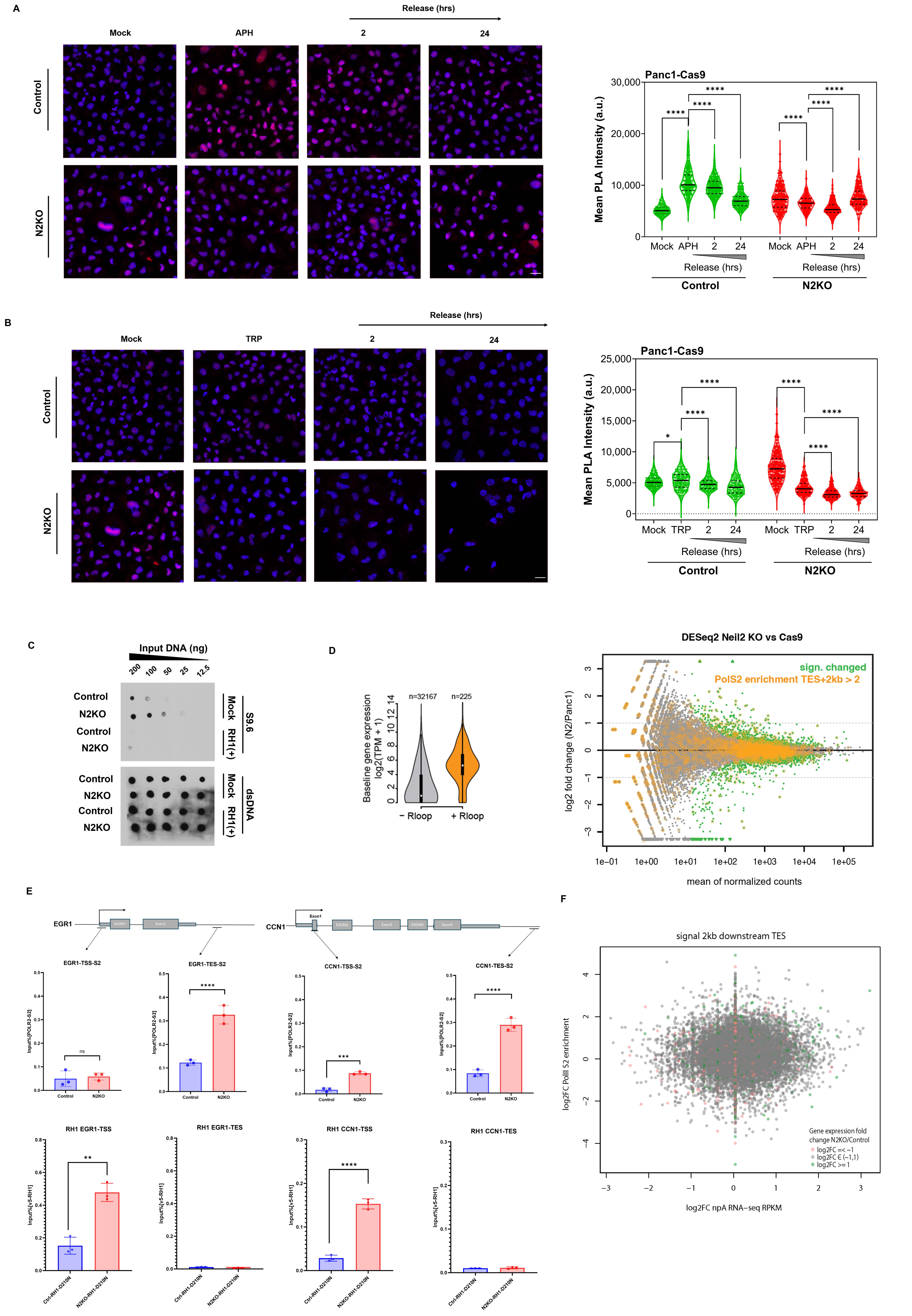
A
Release (hrs)
Mock
APH
2
24
Control
N2KO
Panc1-Cas9
Mean PLA Intensity (a.u.)
30,000
20,000
10,000
0
****
Mock APH 2 24 Mock APH 2 24
Release (hrs)
Control
N2KO
B
Release (hrs)
Mock
TRP
2
24
Control
N2KO
Panc1-Cas9
Mean PLA Intensity (a.u.)
25,000
20,000
15,000
10,000
5,000
0
*
****
Mock TRP 2 24 Mock TRP 2 24
Release (hrs)
Control
N2KO
C
Input DNA (ng)
200
100
50
25
12.5
Control
N2KO
Control
N2KO
Mock
RH1(+)
S9.6
dsDNA
D
Baseline gene expression
log2(TPM + 1)
n=32167
n=225
− Rloop
+ Rloop
DESeq2 Neil2 KO vs Cas9
sign. changed
PolS2 enrichment TES+2kb > 2
log2 fold change (N2/Panc1)
mean of normalized counts
1e−01
1e+00
1e+01
1e+02
1e+03
1e+04
1e+05
E
EGR1
EXON1
Exon2
CCN1
Exon1
EXON2
Exon3
EXON4
Exon5
EGR1-TSS-S2
EGR1-TES-S2
CCN1-TSS-S2
CCN1-TES-S2
Input%[POLR2-S2]
ns
****
***
Control
N2KO
RH1 EGR1-TSS
RH1 EGR1-TES
RH1 CCN1-TSS
RH1 CCN1-TES
Input%[v5-RH1]
**
Ctrl-RH1-D210N
N2KO-RH1-D210N
F
signal 2kb downstream TES
log2FC PolII S2 enrichment
log2FC npA RNA−seq RPKM
Gene expression fold
change N2KO/Control
log2FC =< −1
log2FC ∈ (−1,1)
log2FC >= 1

Fig.S5

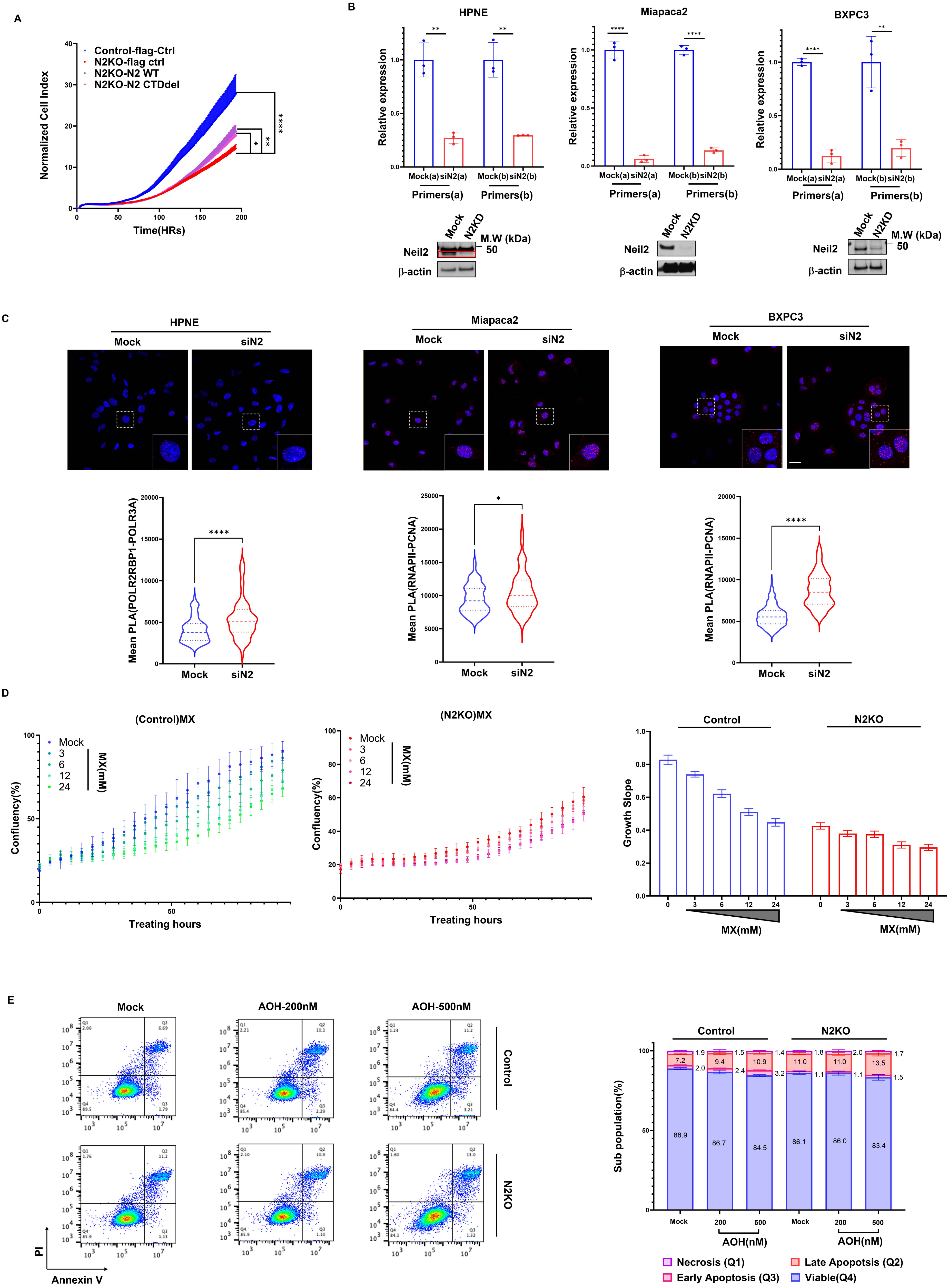
A
Control-flag-Ctrl
N2KO-flag ctrl
N2KO-N2 WT
N2KO-N2 CTDdel
Normalized Cell Index
Time(HRs)
B
HPNE
Miapaca2
BXPC3
Relative expression
Mock(a) siN2(a)
Mock(b) siN2(b)
Primers(a)
Primers(b)
Mock
N2KD
M.W (kDa)
50
Neil2
β-actin
C
HPNE
Miapaca2
BXPC3
Mock
siN2
Mean PLA(POLR2RBP1-POLR3A)
Mean PLA(RNAPII-PCNA)
D
(Control)MX
(N2KO)MX
Mock
3
6
12
24
MX(mM)
Confluency(%)
Treating hours
Control
N2KO
Growth Slope
E
Mock
AOH-200nM
AOH-500nM
Control
N2KO
PI
Annexin V
Sub population(%)
AOH(nM)
Necrosis (Q1)
Late Apoptosis (Q2)
Early Apoptosis (Q3)
Viable(Q4)
